# Loss of the autism associated gene *Tbr1* disrupts prediction and encoding by prefrontal ensembles during socioemotional behaviors

**DOI:** 10.1101/2025.01.21.633988

**Authors:** Marc Turner, Sarah Robinson-Schwartz, Siavash Fazel-Darbandi, John L.R. Rubenstein, Vikaas S. Sohal

## Abstract

Disruptions in many genes linked to autism spectrum disorder (ASD) affect synaptic function and socioemotional behaviors in mice. However, exactly how synaptic dysfunction alters neural activity patterns underlying behavior remains unknown. We addressed this using mice lacking the high confidence ASD gene *Tbr1* in cortical layer 5 (L5) projection neurons (*Tbr1* cKO mice). These mice have known deficits in synaptic input to L5 neurons and social behavior. We also find some abnormalities in anxiety-related avoidance. Calcium imaging of prefrontal L5 neurons revealed that despite reduced overall activity, cKO mice recruit normal numbers of neurons into prefrontal ensembles encoding social and anxiety-related behaviors. However, the stability, inter-neuronal coordination, and reactivation of social ensembles were diminished in cKO mice. Furthermore, in cKO mice, ensembles no longer predicted approach-avoidance decisions. These results reveal new aspects of how prefrontal ensembles encode socioemotional behaviors, and malfunction in the setting of ASD-linked gene disruption.

## INTRODUCTION

Autism spectrum disorder (ASD) encompasses a range of neurodevelopmental conditions defined by specific core features, primarily deficits in the social domain as well as repetitive behaviors. >100 genes have been identified which impart a large individual risk for ASD (Satterstrom et al., 2020; H. R. Willsey et al., 2022). Many of these have been linked to specific cell-types, molecular pathways and cellular processes, and in many cases, mice which model disruptions in these genes have been shown to exhibit altered social behavior. Despite this progress, there remains a critical gap in our understanding between the cellular/molecular and behavioral levels. In particular, how exactly genetic disruptions alter the patterns of neural activity that normally support ASD-relevant behaviors such as social interaction is still largely unknown.

Within the field of ASD genetics, much effort has been made to identify common molecular and cellular pathways impacted by the risk-gene network. Additionally, the temporal and spatial expression of these risk genes have been leveraged to pinpoint convergent brain areas and developmental timepoints that may be particularly critical for ASD pathology (Satterstrom et al., 2020; A. J. Willsey et al., 2013). These efforts have highlighted deep layer projection neurons in prefrontal cortex (PFC) during midfetal development as a spatial and temporal locus for risk-gene convergence. From the ASD risk-gene network, T-box, brain, 1 (*TBR1*) was found to be the most connected to other risk-genes in the PFC at this critical developmental stage (A. J. Willsey et al., 2013).

*TBR1* is a transcription factor expressed in most immature excitatory neurons in layers 5 and 6 of the frontal and motor cortices, whereas layer 5 expression in other cortical regions is reduced (Bulfone et al., 1995; Fazel Darbandi et al., 2020). Previous studies have established that *TBR1* plays a critical role in maintaining the cell identity of layer 6 neurons (Bedogni et al., 2010; Fazel Darbandi et al., 2018; Hevner et al., 2001; McKenna et al., 2011), binds directly to the promoters of many other ASD risk-genes (Fazel Darbandi et al., 2020; Notwell et al., 2016), and exhibits functional convergence with other ASD-associated transcriptional regulators (Darbandi et al., 2024). We recently published a study of mice with the conditional, postnatal deletion of *Tbr1* from cortical layer 5 (L5) neurons. These L5 cKO mice are generated using a Cre-driver line (*Rbp4*-Cre) which leads to deletion at murine post-natal day 0 (P0), which roughly corresponds to human midfetal development (Workman et al., 2013). These conditional knockout (cKO) mice spend less time than their wild-type littermates interacting with novel conspecifics (Fazel Darbandi et al., 2020). Notably, in these cKO mice, both anatomical and physiological measures of excitatory and inhibitory synaptic inputs to L5 neurons are reduced, and spines on L5 neurons have a persistently immature, filamentous morphology. These phenotypes can be rescued by increasing WNT-signaling via genetic interventions, or through lithium treatment (Fazel Darbandi et al., 2020, 2022).

Despite this detailed information on cellular, molecular and behavioral phenotypes in cKO mice, it remains unknown how this genetic disruption affects the patterns of neural circuit activity which mediate behavior. In fact, there is limited information about how the activity of neurons within the prefrontal cortex is altered during behavior in mouse models of autism.

Previous work from our lab has shown that social behavior recruits excessive activity in medial PFC (mPFC) neurons in *Shank3* knockout (KO) mice (Frost et al., 2021). This results in a reduction in the ability of coordinated multineuron activity to specifically encode social interaction. Other work in *Cntnap2* KO mice has found increased baseline activity but decreased recruitment of mPFC neurons by social odors (Levy et al., 2019). Together with more variable neuronal activity, this impairs the encoding of whether odor stimuli belong to social vs. nonsocial categories. Whether these types of abnormalities are present in other mice with genetic disruptions related to autism, e.g., *Tbr1* cKO mice, is unknown. It also remains unknown whether these sorts of abnormalities in multineuron encoding are specific to social behavior, or would also generalize to other prefrontal-dependent behaviors.

This study sought to address these questions using microendoscopic calcium (Ca2+) imaging in freely behaving *Tbr1* cKO mice. Notably, we also find that under some conditions, these mice may exhibit abnormal anxiety-related avoidance, which, like social interaction, depends on the mPFC (Adhikari et al., 2011). This allows us to directly test whether similar abnormalities in neural circuit activity occur during both behaviors. Furthermore, given that a single dose of lithium rescues both synaptic and social interaction deficits in *Tbr1* cKO mice (Fazel Darbandi et al., 2020), we could examine whether this treatment elicits any long-lasting normalizations of *in vivo* activity.

In contrast to previous findings from *Shank3* KO and *Cntnap2* KO mice, we find that *Tbr1* cKO mice have decreased (not increased) L5 mPFC activity across multiple behaviors. Despite this overall reduced activity, social behavior still recruits normal numbers of mPFC neurons in *Tbr1* cKO mice. However, similar to *Shank3* KO mice, the coordination of neuronal coactivity within social behavior-encoding ensembles is disrupted. Interestingly, this defect in multineuron coordination appears to be specific to social behavior and was not observed during anxiety-related avoidance in the elevated plus maze (EPM). We identified other novel aspects of prefrontal ensemble function related to social behavior (ensemble reactivation following interactions with novel mice) and anxiety-related avoidance (ensemble activity predicts approach vs. avoidance), and find that these are disrupted in *Tbr1* cKO mice. Finally, we find that a single treatment with lithium ameliorates the deficient activity of L5 cKO neurons.

## MATERIALS AND METHODS

### Animal Subjects

All animal care procedures and experiments were conducted in accordance with the National Institutes of Health guidelines and approved by the Administrative Panels on Laboratory Animal Care at the University of California, San Francisco. Mice were housed in a temperature-controlled environment (22–24 °C) with ad libitum access to food and water. Mice were reared in normal lighting conditions (12-h light/dark cycle). *Tbr1* layer 5 knockout mice were generated by crossing *Tbr1^f/+^* with *Tbr1^f/+^::Rpb4-cre^±^::Ai14 ^+/+^* mice (Fazel Darbandi et al., 2020). Experimental mice were either *Tbr1^+/+^::Rpb4-cre^±^::Ai14 ^+/ -^ (Tbr1* WT) or *Tbr1^f/f^::Rpb4- cre^±^::Ai14 ^+/ -^ (Tbr1* cKO).

### Surgeries

All animals used for microendoscopic recordings underwent two surgeries: the first for viral injection and the second for GRIN lens implantation. For both surgeries, mice were anaesthetized with isoflurane (3.5% induction, 1-2% maintenance in 95% oxygen, flow rate 0.9L/min) and secured with ear bars in a stereotaxic frame (David Kopf instruments). Analgesics were administered via subcutaneous injection at surgery onset (buprenorphine, 0.1 mg/kg; meloxicam, 2 mg/kg). Body temperature was maintained using a heating pad. The scalp was incised along the rostro-caudal axis to expose the dorsal surface of the skull, which was then aligned using bregma and lambda as references. After surgery, animals were allowed to recover on a heated pad until ambulatory. All animals received an additional dose of meloxicam on the day following surgery.

Viral injections were performed on mice 10-12 weeks old. A dental drill was used to create an opening in the skull above the right mPFC (-1.7mm AP, +0.4mm ML relative to bregma). AAV9-hSyn-FLEX-jGCaMP7f-WPRE (AddGene) was diluted 3:1 in sterile saline immediately before injection. Virus was injected at four different depths, starting with the most ventral, (-2.75mm, -2.5mm, -2.25mm, and -2.0mm DV; 150nl at each depth) at a rate of 100nl/min with a 35-gauge microinjection syringe (World Precision Instruments) connected to a pump (UMP3 UltraMicroPump, World Precision Instruments). The needle was kept at each injection site for 5mins after infusion. Scalp incisions were sutured.

GRIN lens implant surgeries were performed two weeks after viral injection. A dental drill was used to create an 1mm diameter opening in the skull above the right mPFC (-1.7mm AP, +0.55mm ML relative to bregma). The cortex was aspirated to an approximate depth of 2.0mm, and a 1 mm diameter x 4 mm long integrated GRIN lens (Inscopix) was slowly lowered to a final depth of 2.3mm. Implants were secured to the skull with Metabond (Parkell). After surgery, mice were singly housed to prevent damage to their implants. Mice were allowed 3 weeks to recover from surgery before behavioral testing and microendoscopic imaging.

### Behavioral Assays

For all cohorts, mice were habituated to experimenter handling for 3-5 days (5-10min per day). Mice were transported to the testing room 45-60min prior to test onset. For calcium imaging cohorts, mice were also habituated to the head-mounted microscope before testing (3-5 days, 45min per day). Behavioral videos were recorded with a video camera, and real-time animal tracking was performed using AnyMaze software.

### Social Behavior Assays

For the initial behavioral cohort (**Fig. 1A-B**), group-housed animals were transferred to a clean holding cage 1 hour prior to testing. Animals were then, one-by-one, transferred to their home cage for testing. Mice were presented with a novel juvenile conspecific (sex-matched, 4-6 weeks old) for 5min, and then with a novel object (a clean 15mL Falcon tube) for 5min.

**Figure 1.**
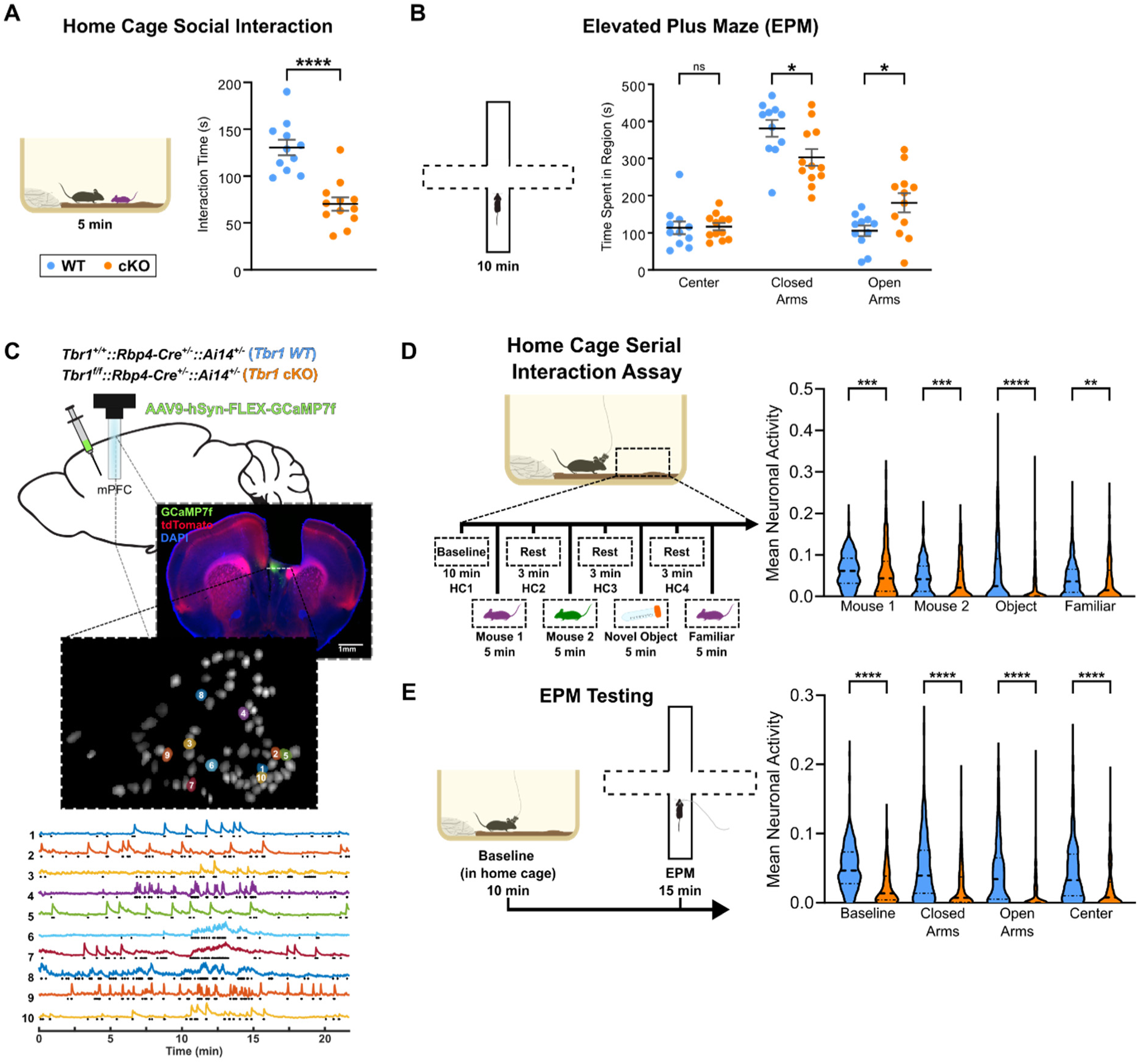
Tbr1 cKO mice exhibit abnormal socioemotional behavior and reduced activity of L5 mPFC neurons. (A) Interaction time with a novel juvenile conspecific during a 5-minute assay in the home cage (n= 23 mice: 11 WT, 12 Tbr1 cKO). (B) Zone time during a 10-minute EPM test (n= 23 mice: 11 WT, 12 Tbr1 cKO). (C) Experimental approach for recording mPFC L5 neurons during behavior. (D) Activity of mPFC L5 neurons during the serial interaction assay. *Left*, timeline of home cage serial interaction assay. *Right,* violin plots of activity of all recorded neurons (n=306 WT, n=278 cKO). Mean activity of each neuron was calculated as the fraction of frames corresponding to active social exploration that had a detected calcium event. P-values shown are from a Kruskal-Wallis test between genotypes. (E) Activity of mPFC L5 neurons during the EPM assay. *Left*, timeline EPM assay. *Right,* violin plots of activity of all recorded neurons (n=325 WT, n=197 cKO). Mean activity of each neuron was calculated as the fraction of frames corresponding to exploration of each EPM zone that had a detected calcium event. P-values shown are from a Kruskal-Wallis test between genotypes.

### Interaction time with both targets was scored manually

For calcium imaging cohorts, mice (singly housed) underwent a serial interaction assay in their home cage. Mice were attached to the head-mounted microscope and an initial 10- minute baseline recording was performed. Following the baseline, four 5-minute interaction epochs were interleaved with 3-minute rest periods (with no interaction target). The four interaction epochs were (in order): a novel same-sex juvenile mouse (Mouse 1), a second novel same-sex juvenile mouse (Mouse 2), a 15mL plastic conical tube (Novel Object), and the first novel juvenile (Mouse 1, now labeled ‘Familiar’). All interactions were manually scored.

### Elevated Plus Maze

For the initial behavioral cohort (**Fig. 1A-B**), animals were placed into the center of the maze and allowed to freely explore for 10min. For calcium imaging cohorts, two days after social testing, animals were attached to the head-mounted microscope and an initial 10-minute baseline recording was performed in the home cage. Mice were then placed into one of the closed arms of the maze and allowed to freely explore for 15min. For all cohorts, AnyMaze tracking was used for subsequent behavioral analyses.

### Lithium Chloride Treatment

For the calcium imaging cohorts, a subset of animals were retested on both the serial interaction assay and the EPM after receiving a single IP injection of lithium-chloride (5mg/kg). Animals underwent initial social testing on day 1, initial EPM testing on day 3, and received LiCl on day four. Retesting on these assays was conducted 30 days after LiCl administration.

### Data acquisition and processing

Calcium signals were imaged at 20Hz with 4x spatial downsampling using a head-mounted, one-photon microendoscope (nVoke2; Inscopix Inc.). Behavioral videos were recorded using AnyMaze software, and a TTL cable connecting the AnyMaze computer to the Inscopix DAQ box was used to align calcium activity and behavior videos. Calcium activity videos were preprocessed in Inscopix Data Processing Software (IDPS). Videos were spatially band-pass filtered with cut-offs set to 0.015 pixel-1 (low) and 0.300 pixel-1 (high), and then motion corrected. Preprocessed videos were exported from IDPS as tiff files and imported to MatLab for cell segmentation. Putative cellular ROIs were identified using a combined PCA/ICA approach (Mukamel et al., 2009). For each movie, putative neurons were manually sorted to remove ROIs with low SNR or overlapping boundaries. Cell sorting was done with the assistance of the signalSorter GUI from an open-source calcium imaging software package (CIAPKG) (Corder et al., 2019).

Cellular dF/F traces for sorted neurons were extracted and then used to generate a binary calcium event raster for each neuron using a previously described event detection algorithm (Frost et al., 2021). All subsequent analyses were done on binary event rasters. Briefly, for each dF/F trace, the median deviation from the mean was calculated for each 10- frame window across the entire recording. The median of the lower 50^th^ percentile of these values was taken as the baseline variance of the dF/F trace, σ. Events were initially detected by finding increases in dF/F exceeding 2.5σ over 1 second, then only keeping events that also exceeded a 15σ increase over 2 seconds and had a total AUC of at least 250σ.

### Data analysis

All analyses of calcium activity data were performed in MatLab using custom scripts.

### Determining Mean Cell Activity

To calculate a cell’s mean activity during a particular behavior, first, all calcium imaging frames corresponding to that behavior were identified. Mean activity was then computed as the fraction of these behaviorally associated frames with a detected calcium event.

### Correlation Analysis

To assess the pairwise correlations between neurons during a particular behavior, frames corresponding to that behavior were identified and extracted from the full recording. Pairwise correlations were then calculated on this behavior-specific activity matrix using the MatLab corr function. Any correlation with an associated p-value below 0.05 was taken to be significant, and results were reported as the portion of pairwise correlations that were significant.

### Neural Network Classifiers and Identification of Behavioral Ensembles

To identify behaviorally recruited neural ensembles, a neural network classifier (NNC) was utilized (previously detailed in Frost et al., 2021). NNCs were trained to distinguish two different behavioral periods (B1 and B2) based on binary calcium event rasters, e.g., the baseline period vs. interaction with the first novel mouse (Mouse 1), or the baseline period vs. time spent in the open arms. Each NNC contained a hidden layer comprised of 1000 artificial units. Each hidden layer unit received input from a random subset of real neurons (connection probability was set to 0.3). The binary matrix C (dimensions: N real neurons x n hidden units) described connectivity from real neurons to the hidden layer. Hidden layer units were all connected to a single output unit with weights given by the vector w. The output weight vector w was updated during training.

To train a binary NNC, all calcium imaging frames corresponding to B1 and B2 were identified. Frames associated with B1 and B2 were assigned output values of 1 and 0, respectively. For identification of ensembles, frames were not split into separate training/testing sets because the purpose was to train the weight vector w, not to assess performance. Training consisted of 500 passes through the training frames, using a different, random order on each pass. If B1 and B2 had different numbers of associated frames, the larger set was randomly subsampled to match the number of frames in the smaller set; each training pass used a new random subset. Each pass through the data was further capped at 1000 frames to mitigate disparities between animals. For each animal we trained 20 NNCs for each pair of behaviors. After training, the effective weight associated with each real neuron was computed as **C*w**. The resulting effective weight vectors were then averaged across all 20 NNCs, resulting in one effective weight vector for each behavioral comparison for each animal.

Given the labels of 1 and 0 for B1 and B2 frames, respectively, positive effective weights were associated with B1 encoding neurons and negative weights with B2 encoding neurons. To identify neural ensembles for specific behaviors from the effective weights, a threshold of 1 standard deviation (SD) was used. Thus, the ‘B1 ensemble’ was defined as neurons with an effective weight at least 1 SD above the average value, and the ‘B2 ensemble’ was defined as neurons with an effective weight at least 1 SD below the mean.

For the serial interaction assay data, 3 types of NNC were trained per animal. In all cases, B1 was designated as the baseline home cage recording period, while B2 was either Mouse 1 interaction, Mouse 2 interaction, or Familiar interaction. Only frames corresponding to active social interaction were used.

For the EPM assay data, 2 types of NNC were trained per animal. In both cases, B1 was designated as the baseline home cage recording period, while B2 was either Closed Arm exploration or Open Arm exploration.

### SHARC and Swap Shuffling

SHARC is a method for generating surrogate datasets from an original matrix of neural activity that nonrandomly shuffles blocks of activity such that the shuffled data conforms to a target correlation matrix (Luongo et al., 2016). Thus, the resultant surrogate datasets preserve the pairwise correlation structure of the target correlation matrix that is provided. A modification of this method that also preserves the overall activity of each neuron was used (Frost et al., 2021).

SHARC surrogate datasets were generated for a particular behavior by first creating a subraster consisting only of frames corresponding to that behavior. A pairwise correlation matrix was calculated from this behavioral subraster and used as the target correlation matrix for SHARC in order to preserve the native correlation structure.

Swap-shuffled surrogate datasets were also generated from behavioral subrasters. This method involves making random, reciprocal swaps of detected calcium events between neurons. Resulting swap-shuffled surrogate datasets preserve the overall activity of each neuron and the total network activity of each frame in the native dataset, but the native correlation structure is broken.

To assess the relative contribution of neuronal coactivity in encoding of behavior, 20 SHARC and 20 Swap-shuffled surrogate datasets were generated for each of the behaviors that NNCs were trained on (Social baseline, Mouse 1, Mouse 2, Familiar, EPM baseline, Closed Arm exploration, and Open Arm exploration). These NNCs trained on real data (20 for each behavioral comparison) were then tested on corresponding SHARC and swap-shuffled datasets. For example, the 20 NNCs trained on Social baseline and Mouse 1 data were each tested on one SHARC dataset composed of Baseline and Mouse 1 data, and one swap-shuffled dataset composed of Baseline and Mouse 1 data. Because there were always more Baseline frames than frames for the other behavior, the Baseline surrogate data was randomly subsampled to provide an equal number of frames. For each NNC, a performance improvement score was calculated as (P_SHARC_ - P_Swap_) / (P_Swap_ - 0.5), where P_SHARC_ is performance on SHARC data, P_Swap_ is performance on swap-shuffled data, and 0.5 is the chance performance level. The 20 different improvement scores were averaged to yield a single value for each behavioral comparison in each animal.

### Quantification and statistical analysis

All statistical calculations were performed in MatLab or Graphpad Prism. All details of these statistical analyses are provided in the main text and figure legends.

## RESULTS

### Selective deletion of *Tbr1* from layer 5 cortical neurons causes abnormal social and anxiety-related behaviors

To determine whether *Tbr1* expression in cortical L5 pyramidal neurons is required for normal socioemotional behavior, we crossed the *Rbp4-Cre* mouse line with a line harboring a floxed *Tbr1* allele, as described previously (Fazel Darbandi et al., 2020). This results in *Tbr1* deletion within L5 excitatory neurons of the neocortex at postnatal day zero (P0), ∼8 days after *Tbr1* is first expressed. We refer to these *Tbr1*^fl/fl^::*Rbp4-Cre*^±^::*Ai14*^±^ mice as ‘*Tbr1* cKO’ mice, or just ’cKO.’

Previously, we showed that *Tbr1* cKO mice exhibit deficient social interaction as measured by a home-cage resident-intruder assay with a novel, same-sex juvenile conspecific (Fazel Darbandi et al., 2020). To determine if these mice display additional behavioral deficits, cKO mice and WT littermates (*Tbr1*^+/+^::*Rbp4-Cre*^±^::*Ai14*^±^) underwent a battery of behavioral tests. As seen previously, cKO mice displayed abnormal social behavior as measured by interaction time with a novel same-sex juvenile mouse in the home cage (WT: 131 ± 28 sec, cKO: 70 ± 24s, p = 0.000020, Welch’s Unpaired T-test) (Fig. 1A). By contrast, there was no difference in the time WT vs. cKO mice spent exploring a novel object (data not shown).

Intriguingly, *Tbr1* cKO mice also had a deficit in anxiety-related avoidance as measured using the elevated plus maze (EPM). cKO mice spent significantly more time exploring the open arms (181 ± 90 sec in cKO vs. 106 ± 47 sec for WT, p = 0.027, 2-way ANOVA with Sidaks correction) and significantly less time in the closed arms (303 ± 79 sec in cKO vs. 381 ± 56 sec for WT, p = 0.020, 2-way ANOVA with Sidak’s correction) (Fig. 1B). In the open field test (OFT), another commonly-used assay of anxiety-related avoidance, there were no significant differences in the amount of time spent in the center vs. the perimeter zones. However, cKO mice did make significantly more entries into the center zone, despite displaying comparable levels of locomotion with WT mice (Fig. S1A-C). While the EPM and OFT are not completely interchangeable assays, this increase in entries into and distance traveled within the OFT center aligns with the decreased open arm avoidance in the EPM, supporting the interpretation that *Tbr1* cKO mice may, under certain conditions, exhibit decreased anxiety-related avoidance behaviors.

### *Tbr1* cKO neurons have significantly reduced activity during behavior

To better understand how the loss of *Tbr1* affects neuronal circuits underlying social and anxiety-related behaviors, we used miniaturized, head-mounted microscopes to record Ca2+ signals from L5 mPFC neurons in freely moving cKO mice and their WT littermates. Mice were injected at 11-13 weeks of age with a Cre-dependent virus encoding the fluorescent calcium indicator GCaMP7f (AAV9-hSyn-FLEX-GCaMP7f-WPRE). Thus, we recorded signals specifically from L5 neurons that were labeled by the Rbp4-Cre line, i.e., the neurons from which *Tbr1* had been deleted (Fig. 1C). Mice were implanted with a 1mm gradient refractive index (GRIN) lens over mPFC 2-3 weeks after virus injection, then allowed to recover for 3 weeks before habituation to the microscope and behavioral testing.

Mice were first tested on a home cage social interaction assay, which consisted of an initial 10-minute baseline recording period followed by four 5-minute interaction epochs interleaved with 3-minute rest periods. The four interaction epochs were (in order): a novel same-sex juvenile mouse (Mouse 1), a second novel same-sex juvenile mouse (Mouse 2), a 15mL plastic conical tube (Novel Object), and finally a re-exposure to the first novel juvenile (Mouse 1, now labeled ‘Familiar’) (Fig. 1D). Two days later, mice had a 10-minute baseline recording in their home cage, followed by 15 minutes of testing in the EPM (Fig. 1E).

Calcium signals were imaged at 20Hz using the Inscopix nVoke system. Following video preprocessing, putative cellular ROIs were identified using a combined PCA/ICA approach and manually sorted (Mukamel et al., 2009). Cellular dF/F traces were extracted and a custom event detection algorithm was used to obtain a binary calcium-event raster for each neuron (Frost et al., 2021) (Methods). All subsequent analyses were performed using these binary event rasters.

First, to determine how the loss of *Tbr1* affected the activity of L5 neurons, we calculated the mean activity of each neuron for relevant timepoints within both the serial interaction assay and the EPM. Mean activity was calculated for each neuron as the fraction of frames with a detected calcium event out of all frames corresponding to a given behavior. For the interaction assay, only frames corresponding to “active interaction” were used. “Active interaction” included all instances in which the test mouse was within ∼1cm of, and oriented toward, the target (either a juvenile mouse or novel object). Despite similar activity levels of WT and cKO neurons during the initial baseline period (WT: 0.0399 ± 0.0015, cKO: 0.0422 ± 0.0020; Fig. S1G), during the Mouse 1 (WT: 0.0627 ± 0.0022, cKO: 0.0557 ± 0.0034), Mouse 2 (WT: 0.0461 ± 0.0022, cKO: 0.0372 ± 0.0027), Novel Object (WT: 0.0570 ± 0.0045, cKO: 0.0275 ± 0.0041), and Familiar (WT: 0.0443 ± 0.0024, cKO: 0.0388 ± 0.0031) interaction epochs, cKO neurons were overall significantly less active than their WT counterparts (Fig. 1D). Similarly, in the EPM, cKO neurons were significantly less active in every zone (closed arms, WT: 0.0503 ± 0.0025, cKO: 0.0237 ± 0.0022; open arms, WT: 0.0442 ± 0.0025, cKO: 0.0185 ± 0.0024; center, WT: 0.0451 ± 0.0024, cKO: 0.0210 ± 0.0021) (Fig. 1E). cKO neurons were also significantly less active in the 10 minute baseline recording immediately prior to the EPM (WT: 0.0520 ± 0.0018, cKO: 0.0236 ± 0.0018), despite the absence of a baseline deficit two days earlier (just before social interaction).

Within our smaller imaging cohort, there were not significant decreases in interaction time for any of the individual social targets (Fig. S1E), though the cumulative interaction time exhibited a trend towards reduction that was grossly similar to that observed in our larger, behavior-only cohort (WT: 508 ± 52, cKO: 347 ± 67; p = 0.087) (Fig. S1F). However, in the EPM, our imaging cohort of cKO animals did not exhibit even a trend towards increased open arm exploration time, suggesting that changes in anxiety-related behaviors are less robust across experimental conditions (portion of time spent in open arms: WT = 0.283 ± 0.072, cKO = 0.141 ± 0.035, p = 0.33) (Fig. S1D).

### A neural network classifier approach for identifying behaviorally recruited neuronal ensembles

To determine whether this reduction in L5 neuronal activity affected the encoding of behavior, we used a neural network classifier (NNC) to identify behaviorally relevant neuronal ensembles (**Fig. 2A**). Previous work from our lab showed that this approach can identify both neurons which encode aspects of behavior via changes in their individual activity levels and groups of neurons which encode this information collectively, e.g., via increases or decreases in their correlated activity (coactivity) during specific behaviors (Frost et al., 2021).

**Figure 2.**
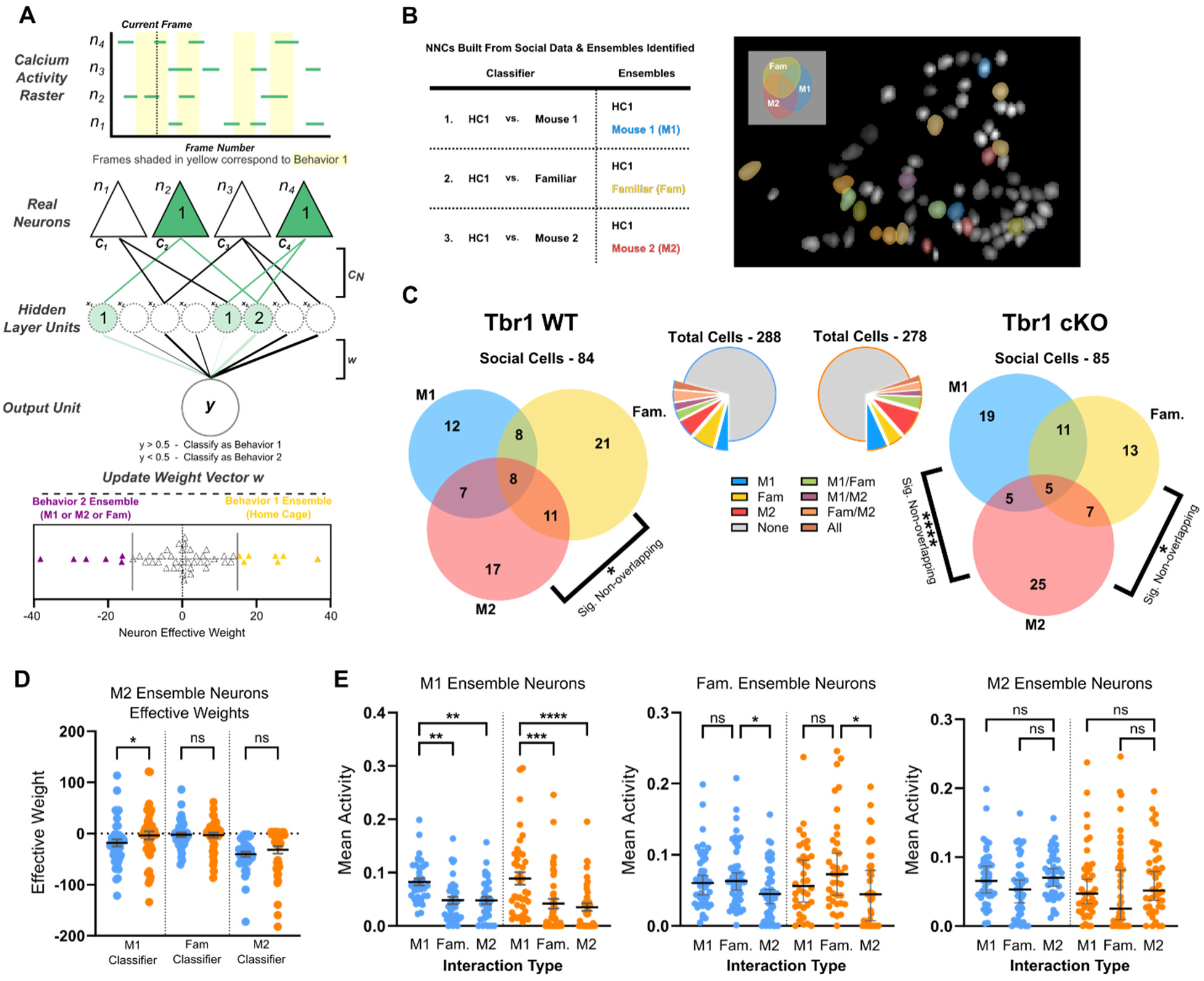
Neural ensembles recruited during social behavior are more stable across different interactions in WT than in cKO mice. (A) Overview of NNC architecture and schematic showing how neuronal ensembles encoding interactions with mouse 1 (‘M1’), mouse 2 (‘M2’), or mouse 1 when it was presented a second time (‘Fam’) were identified based on the weights of connections in trained NNCs. (B) *Left,* summary of the different NNCs that were trained for each mouse, and the ensembles identified using each NNC. *Right,* example FOV from one mouse in which the colors of different neurons indicate their membership in one or more of the three social ensembles (blue, red, yellow = M1, M2, or familiar ensembles, respectively; purple, orange and green = overlapping membership in two ensembles). (C) Summary of identified social cells. *Middle*, pie charts indicating the fraction of recorded cells identified as socially encoding for WT (left, blue border) and cKO (right, orange border) mice (n=10 mice: 5 WT, 5 cKO). *Left,* Venn diagram showing overlaps between WT social ensembles. *Right,* Venn diagram displaying overlaps of cKO social ensembles. In both groups, we found significantly less-than-chance overlap between the M2 and Fam ensembles. In cKO animals, there was also significantly less-than-chance overlap between the M1 and M2 ensembles. Significance of overlap determined by one-tailed Fisher’s Exact tests, performed separately for each genotype and pair of ensembles. (D) Effective weights of M2 neurons (n = 43 for WT, n=42 for cKO) in each of the three trained NNCs from (B). M2 neuron weights in the M1 classifier are significantly shifted away from zero (towards negative values) in WT mice, compared to cKO. This reflects the fact that in WT but not cKO mice, neurons in the M2 ensemble also tend to encode interactions with M1. P-values shown are from Kruskal-Wallis tests with Dunn’s multiple comparison correction between genotypes. (E) Activity of neurons in each of the three social ensembles during each type of interaction. During social interactions, activity in social ensembles was recruited similarly in both genotypes. *Left,* M1 neurons (n = 35 and 40 for WT and cKO, resp.). *Middle*, Fam. neurons (n = 48 and 36 for WT and cKO, resp.). *Right*, M2 neurons (n = 43 and 42 for WT and cKO, resp.). P-values shown are from Kruskal-Wallis tests with Dunn’s multiple comparison correction within genotypes.

For each animal, NNCs were trained to distinguish two different behavioral periods (B1 and B2) based on calcium event data. For example, B1 could correspond to frames during the baseline home cage period (when the mouse is alone), whereas B2 could be frames corresponding to periods of active social interaction. Each NNC contained a hidden layer comprised of 1000 artificial units, each of which received input from a random subset of real neurons based on a connection probability of 0.3. The binary matrix **C** (dimensions: *N* real neurons x 1000 hidden units) described connectivity from real neurons to the hidden layer.

Hidden layer units were all connected to a single output unit. The vector **w**, containing the weights of the output connections, was updated during training. After training, the effective weight associated with each real neuron, i.e., the degree to which each neuron tends to increase or decrease the activity of the output unit in order to bias the classification in one direction or the other, could be obtained by calculating the matrix product **C**\***w**.

During training, frames associated with B1 and B2 were assigned output values of 1 and 0, respectively. This means neurons encoding B1 and B2 correspond to those neurons with the largest (most positive) and smallest (most negative) effective weights, respectively. To identify neural ensembles encoding specific behaviors, we used a threshold of 1 standard deviation (SD). I.e., the ‘B1 ensemble’ was defined as neurons with an effective weight at least 1 SD above the average value, and the ‘B2 ensemble’ was neurons with an effective weight at least 1 SD below the average value.

The major advantage of defining neuronal ensembles in this manner, versus simply identifying which neurons are most active during a particular behavior, is that NNCs can also identify neurons that encode behavior collectively, via changes in their correlations or coactivity. By contrast, defining ensembles based simply on the activity of each neuron would overlook the potential for this type of inter-neuronal synergy or combinatorial coding. Our previous work showed that optimal linear decoders based on logistic regression or support vector machines are also insensitive to coactivity-based encoding (Frost et al., 2021).

#### Ensembles encoding different conspecifics become abnormally differentiated in *Tbr1* cKO mice

To test for altered social encoding in *Tbr1* cKO, we trained three types of NNC for each animal, so as to identify social ensembles corresponding to three types of social interaction: ‘M1’ (interaction with Mouse 1 when it is novel), ‘Fam’ (interaction with Mouse 1 when it is familiar), and ‘M2’ (interaction with Mouse 2). To identify each of these ensembles, a NNC was trained using data from the baseline recording period (HC1) (Behavior 1) and the relevant social interaction period (Behavior 2) (**Fig. 2B**).

Overall, the fractions of neurons identified as being part of each social ensemble were similar across mice and genotypes. ‘M1’, ‘M2’, and ‘Fam’ neurons were pooled by genotype across mice (n = 84 social cells for WT; n = 85 social cells for cKO), and the overlap between different ensembles was assessed (**Fig. 2C**). In both WT and cKO mice, the Fam and M2 ensembles were significantly non-overlapping, i.e., the number of neurons in both ensembles was significantly less than would be expected by chance (WT: 24/84 in M2 only, 29/84 in Fam only, 19/84 in both, 12/84 in neither, p = 0.017; cKO: 30/85 in M2 only, 24/85 in Fam only, 12/85 in both, 19/85 in neither, p = 0.0098, one-tailed Fisher’s exact test). Interestingly, in cKO mice, the M1 and M2 ensembles were significantly non-overlapping, whereas in WT mice these ensembles overlapped as expected by chance (cKO: 32/85 in M2 only, 30/85 in M1 only, 10/85 in both, 13/85 in neither, p = <0.0001; WT: 28/84 in M2 only, 20/84 in M1 only, 15/84 in both, 21/84 in neither, p = 0.2686; one-tailed Fisher’s exact test) (**Fig. 2C**). This suggests that even though Mouse 1 and Mouse 2 interactions have something in common – both involve a novel conspecific – their encoding becomes abnormally distinct in cKO mice.

To further test the idea that the encoding of Mouse 1 and Mouse 2 becomes abnormally distinct in cKO mice, we can also examine the pattern of effective weights. Each effective weight represents the degree to which the corresponding neuron encodes one behavior vs. the other, and there is a different set of effective weights associated with each classifier. Thus, we can take an ensemble identified from one classifier (i.e., the ‘M2 ensemble’ derived from classifier which detects interactions with Mouse 2) and look at the effective weights of those neurons in other classifiers to assess the degree to which this ensemble also encodes other behaviors.

Doing this for M2 ensemble neurons, we see that their effective weights in the Mouse 1 classifier are shifted toward negative values in WT mice, indicating that the M2 ensemble also tends to encode Mouse 1 interaction (**Fig. 2D**). However, in cKO mice, effective weights of M2 neurons in the Mouse 1 classifier are significantly shifted towards zero, implying that in cKO animals, the M2 ensemble does not encode Mouse 1 interaction as strongly as in WT animals (average effective weight of M2 neurons in the Mouse 1 classifier = -18.0 ± 6.70 in WT vs. -3.53 ± 7.91 in cKO, p = 0.0420, Kruskal-Wallis test with Dunn’s correction). Thus, the encoding of different novel conspecifics becomes less stable in *Tbr1* cKO mice compared to WT animals.

To more generally examine overall activity levels within each neuronal ensemble and across behaviors, we plotted the average activity of neurons in the M1, M2 or Fam ensembles during Mouse 1, Mouse 2, and Familiar interactions (**Fig. 2E**). Notably, despite the marked reduction in overall activity in cKO mice, activity levels of social ensemble neurons during social behavior were very similar in WT and cKO mice. The modulation of this activity during different types of social behavior was also similar across genotypes. For example, in both WT and cKO mice, the M1 ensemble was significantly more active during Mouse 1 interaction than during Mouse 2 or Familiar interactions, and the Fam ensemble was significantly more active during Familiar interaction than during Mouse 2. Thus, focusing on neuronal ensembles reveals that even though overall population activity during behavior is reduced in cKO mice, activity levels within socially-encoding ensembles are grossly intact. However, their composition become more unstable / variable across interactions with different conspecifics in *Tbr1* cKO mice.

### Social ensemble reactivation following social interactions is lost in *Tbr1* cKO mice

Previous studies have reported the presence of “social memory” neurons in the mPFC (Xing et al., 2021), and there is a rich literature on the importance of neuronal reactivation in promoting memory formation and consolidation (Kaefer et al., 2020; J.-H. Lee et al., 2023; Nishimura et al., 2021; Oliva et al., 2020; Preston & Eichenbaum, 2013). Moreover, several studies have reported social memory deficits in ASD mouse models (Bertoni et al., 2021; Niu et al., 2024; Tao et al., 2022). Therefore, we investigated whether social ensembles might reactivate during the rest periods between interaction epochs. Interestingly, in WT mice we observed a significant elevation of activity in the WT M1 ensemble neurons specifically during the HC2 rest period, which immediately follows Mouse 1 interaction (mean M1 ensemble activity during HC1: 0.0210 ± 0.0026; during HC2: 0.0387 ± 0.0060; during HC3: 0.0260 ± 0.0039; during HC4: 0.0241 ± 0.0035; p = 0.0016, 0.021, 0.028 for HC2 vs. HC1, HC3, HC4, respectively; **Fig. 3A**). By contrast, no such reactivation occurred for Tbr1 cKO M1 neurons (mean M1 ensemble activity during HC1: 0.0159 ± 0.0028; during HC2: 0.0171 ± 0.0040; during HC3: 0.0136 ± 0.0027; during HC4: 0.0137 ± 0.0029; p = 0.97, 0.69, 0.75 for HC2 vs. HC1, HC3, HC4) (**Fig. 3B**). We additionally confirmed that when the activity of M1 ensemble neurons is plotted for HC2 vs. for HC1, the slope of the best fit line is significantly larger for WT than cKO neurons, further illustrating the discordance in reactivation between genotypes (p = 0.0257, F = 5.189, DFn = 1, DFd = 71; **Fig. 3C**). To determine if reactivation is unique to socially-encoding cells, we compared the activity of HC1 ensemble neurons across rest epochs. In both genotypes, the activity dynamics of HC1 ensemble neurons are similar, with activity peaking during the HC1 epoch then decreasing in later rest epochs (**Fig. 3D-F)**.

**Figure 3.**
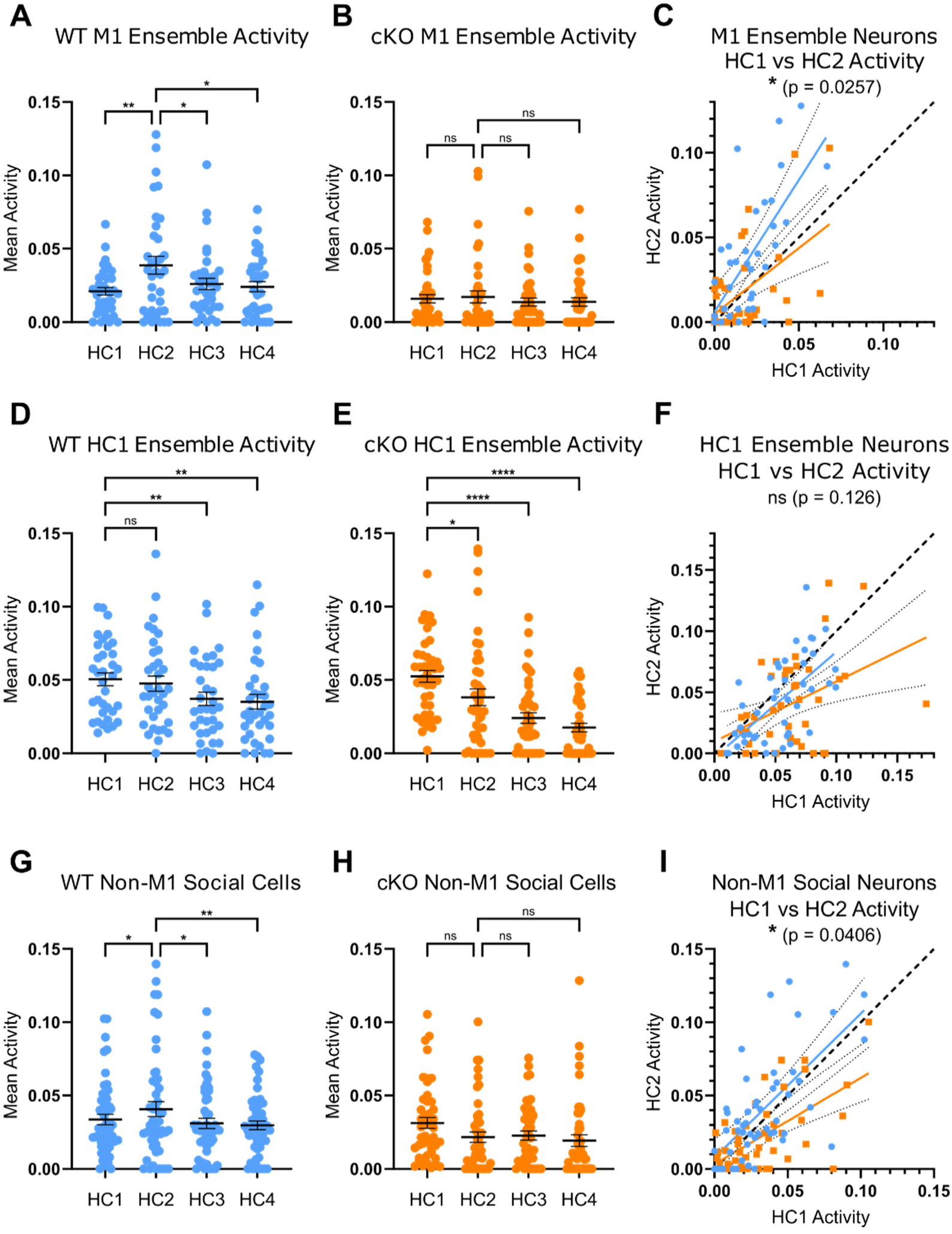
Social neurons reactivate in Tbr1 WT but no cKO animals. (A) Activity of neurons in the WT M1 ensemble (n = 35) during rest epochs of the serial interaction assay. P-values shown are from a RM one-way ANOVA with Dunnett’s multiple comparison test, comparing activity during each rest epoch to activity during the HC2 epoch. (B) Activity of neurons in the cKO M1 ensemble (n = 40) during rest epochs of the serial interaction assay. P-values shown are from a RM one-way ANOVA with Dunnett’s multiple comparison test, comparing activity during each rest epoch to activity during the HC2 epoch. (C) Scatter plot displaying the relationship between HC1 and HC2 activity of WT (blue) and cKO (orange) M1 neurons. The best fit lines from simple linear regression analyses have significantly different slopes, indicating stronger reactivation in WT. (D) Activity of neurons in the WT baseline (BL) ensemble (n = 35) during rest epochs of the serial interaction assay. P-values shown are from a RM one-way ANOVA with Dunnett’s multiple comparison test, comparing activity during each rest epoch to the activity during the HC2 epoch. (E) Activity of neurons in the cKO BL ensemble (n = 43) during rest epochs of the serial interaction assay. P-values shown are from a RM one-way ANOVA with Dunnett’s multiple comparison test, comparing activity during each rest epoch to the activity during the HC2 epoch. (F) Scatter plot displaying the relationship between HC1 and HC2 activity of WT (blue) and cKO (orange) BL neurons. The slopes of the best fit lines from simple linear regression analyses are not significantly different. (G) Activity of WT social neurons that are not in the M1 ensemble (n = 51) during rest epochs of the serial interaction assay. P-values shown are from a RM one-way ANOVA with Dunnett’s multiple comparison test, comparing activity during each rest epoch to the activity during the HC2 epoch. (H) Activity of cKO social neurons that are not in the M1 ensemble (n = 47) during rest epochs of the serial interaction assay. P-values shown are from a RM one-way ANOVA with Dunnett’s multiple comparison test, comparing activity during each rest epoch to the activity during the HC2 epoch. (I) Scatter plot displaying the relationship between HC1 and HC2 activity of WT (blue) and cKO (orange) non-M1 social neurons. The best fit lines from simple linear regression analyses have significantly different slopes, indicating that WT social neurons, even those not in the M1 ensemble, display stronger activation.

We also looked at the activation of non-M1 social neurons (i.e., Fam. and M2 ensemble neurons that did not overlap with the M1 ensemble) during rest epochs. We did not see an analogous elevation in activity of M2 ensemble neurons in the rest period immediately following Mouse 2 interaction (HC3; there was no rest period recording following Familiar interaction) (**Fig. S3**). However, similar to M1 neurons, we observed an elevation of activity during the HC2 epoch compared to HC1 for non-M1 social neurons in WT (mean non-M1 social neuron activity during HC1: 0.0336 ± 0.0035; during HC2: 0.0407 ± 0.0051; during HC3: 0.0310 ± 0.0035; during HC4: 0.0297 ± 0.0029; p = 0.050, 0.015, 0.0074 for HC2 vs. HC1, HC3, HC4 respectively). This increased activity of non-M1 social neurons during HC2 was not present in cKO mice (mean non-M1 social neuron activity during HC1: 0.0314 ± 0.0037; during HC2: 0.0217 ± 0.0036; during HC3: 0.0227 ± 0.0030; during HC4: 0.0193 ± 0.0040; p = 0.060, 0.99, 0.90 for HC2 vs. HC1, HC3, HC4 respectively) (**Fig. 3G-I)**. This ‘preactivation’ suggests that the elevation in activity that we observed during the rest period following an initial social interaction is not limited to cells which strongly encode that social interaction, but generalizes to all social cells, e.g., cells which may subsequently be recruited into other social ensembles. Furthermore, the absence of such reactivation may relate to the decreased stability of social ensembles in cKO mice.

### The encoding of social behavior by correlated activity is diminished in *Tbr1* cKO mice

We previously showed that pairwise neuronal correlations, or neuronal coactivity, in the mPFC encodes information about social behavior, over and above the information transmitted by neuronal activity levels (Frost et al., 2021). Moreover, this contribution of prefrontal correlations was diminished an autism model: *Shank3* KO mice. To assess the contribution of correlated activity to social encoding in *Tbr1* cKO mice, we measured the ability of NNCs trained on our data to classify social behavior in surrogate datasets that had been shuffled to either preserve or disrupt pairwise correlations between neurons (**Fig. 4A**). Specifically, we generated surrogate datasets using either the <u>SH</u>uffling <u>A</u>ctivity to <u>R</u>earrange <u>C</u>orrelations (SHARC) method (Frost et al., 2021; Luongo et al., 2016), which constraints shuffled data to conform to a target correlation matrix (in this case preserving the original correlations), or randomly swapped activity between neurons, which preserves activity levels across neurons and timepoints while disrupting correlations (Methods).

**Figure 4.**
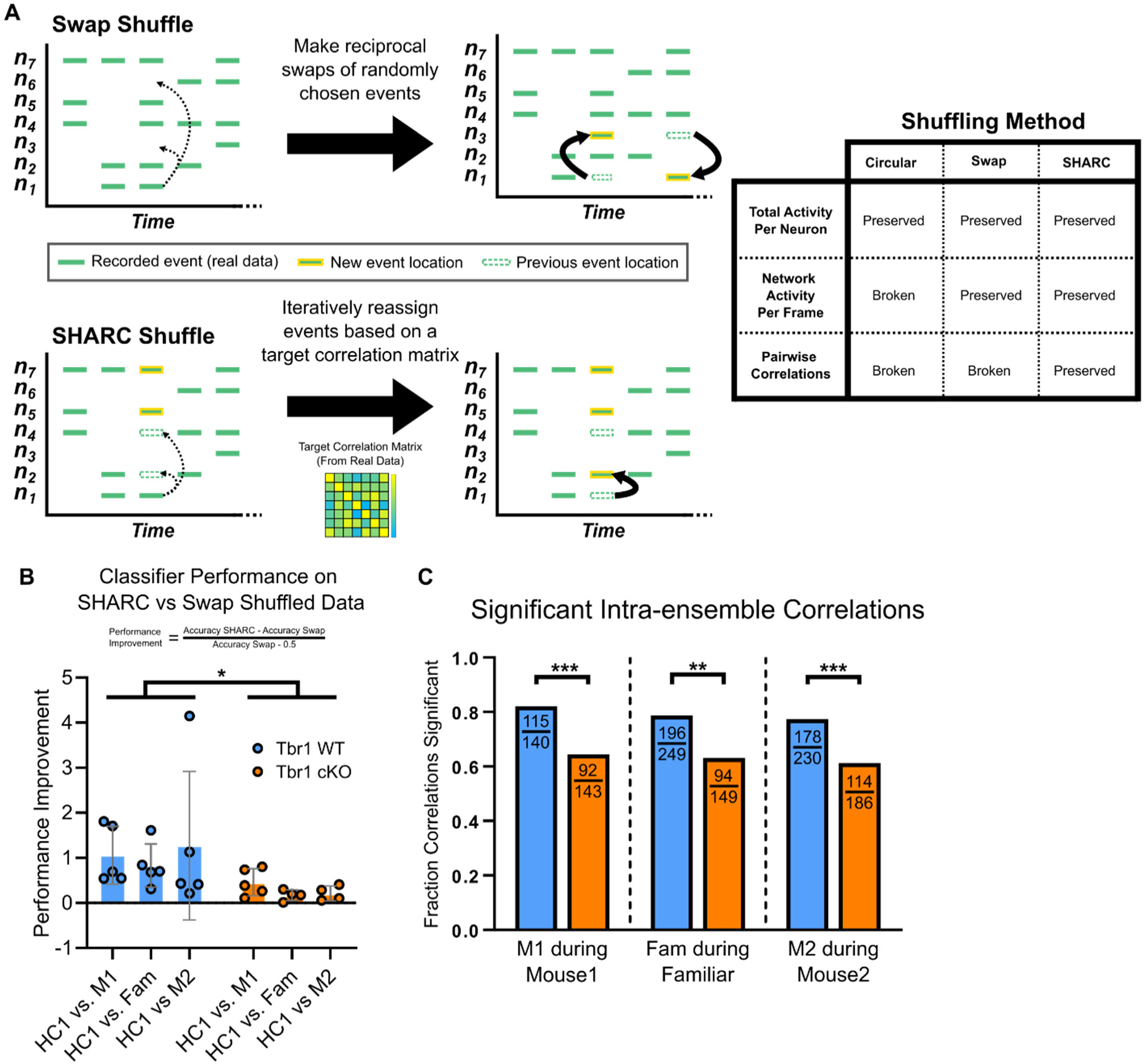
Correlations between neurons contributes more to social encoding in WT than cKO mice. (A) Overview of shuffling methods for binary calcium activity rasters. *Left, top*, schematic of Swap Shuffle method. Random reciprocal swaps of calcium events (blocks of contiguous active frames) between neurons shuffled activity, so as to preserve both the total number of events in each neuron and in each frame. *Left, bottom,* schematic of SHARC method. Calcium events are iteratively reassigned to other neurons based on a target pairwise correlation matrix, preserving total activity in each neuron and each frame, and also preserving correlations present in the original data. *Right*, table summarizing shuffling methods. (B) Performance of classifiers trained on real data and tested on shuffled data. (Note: each subraster corresponding to a specific behavior was shuffled separately). Performance of each trained classifier was assessed on Swap Shuffled and SHARC datasets, and a Performance Improvement score was calculated to assess the contribution of correlated activity in encoding social interaction. A 2-way ANOVA with Sidaks correction of Performance Improvement scores revealed a significant effect of genotype (p = 0.0183). (C) The fraction of significant pairwise correlations within each social ensemble during its corresponding interaction epoch is plotted. Pairwise correlations and their p-values were calculated using the Matlab *corr* function. Correlations with an associated p-value < 0.05 were taken as significant. P-values for genotype differences in the fraction of significant pairwise correlations were calculated using Fisher’s Exact tests for each ensemble.

For each mouse, we identified activity rasters associated with the baseline period or each type of social interaction (Mouse 1, Mouse 2, and Familiar). Then, for each of these rasters we generated 20 SHARC and 20 Swap-shuffled surrogates. The NNCs described previously (**Fig. 2B**), which were trained on the original unshuffled data, were then tested using the corresponding SHARC and Swap surrogate datasets (i.e., the Mouse 1 NNC was tested on shuffled baseline and Mouse 1 datasets) in order to obtain the performance on SHARC (P_SHARC_) or Swap-shuffled (P_Swap_) data. To quantify the degree to which correlations encode information beyond the information encoded by activity levels alone, we calculated: (P_SHARC_ - P_Swap_) / (P_Swap_ - 0.5), where P_SHARC_ = NNC performance on SHARC shuffled data, P_Swap_ = NNC performance on swap-shuffled data, and 0.5 is chance performance level.

Across all three types of social interaction, the improvement in classifier performance achieved when using SHARC shuffling to preserve correlations was higher for WT than *Tbr1* cKO mice (average normalized improvement = 1.05 ± 0.13 in WT vs. 0.28 ± 0.09 in cKO; significant effect of genotype, p = 0.018 by 2-way ANOVA; **Fig. 4B**). Thus, the normal role of prefrontal coactivity in encoding social interactions is diminished in cKO mice. To more directly relate this finding, derived from measuring classifier performance using surrogate datasets, to the social ensembles we had previously identified in real data, we examined *intra-ensemble correlations*. For each pair of neurons within an ensemble, we assessed whether activity of that pair was significantly correlated during periods of social interaction. We found that the proportion of significant intra-ensemble pairwise correlations (PWCs) was consistently higher in WT compared to *Tbr1* cKO mice (M1 ensembles: 115/140 pairs significant in WT vs. 92/143 in cKO, p = 0.0008 by χ2 test; M2 ensembles: 178/230 pairs significant in WT vs. 114/186 in cKO, p = 0.0005 by χ2 test; Fam ensembles: 196/249 pairs significant in WT vs. 94/149 in cKO, p = 0.0011 by χ2 test) (**Fig. 4C**). Thus, intra-ensemble correlations are diminished in cKO, likely explaining why prefrontal coactivity contributes less to social encoding in this genotype. To delve further into the nature of this abnormality in intra-ensemble correlations, we plotted, for both WT and cKO mice, the distributions of significant correlations between neurons in the M1 (**Supplemental Fig. S4D**), M2 (**Supplemental Fig. S4E**), or Familiar (**Supplemental Fig. S4F**) ensembles. For all three ensembles, the significant intra-ensemble correlations comprised both a large number of weak negative correlations and a more broad distribution of positive correlations. Interestingly, the decrease in significant intra-social ensemble correlations in cKO mice was consistently (across all three ensembles) driven mainly by the loss of negative (rather than positive) correlations.

### Prefrontal ensembles predict approach-avoidance decisions in WT but not cKO mice

Having identified these abnormalities in social encoding within *Tbr1* cKO mice, we next turned our attention to anxiety-related behaviors, specifically approach/avoidance in the EPM. Again, we used NNCs to derive behavioral ensembles. For each mouse, two NNCs were trained to distinguish baseline periods when the mouse was in its home cage periods from exploration of either the closed or open arms; the resulting two NNCs were then used to derive closed (CL) and open (OP) arm ensembles, respectively (**Fig. 5A**). Furthermore, as we did earlier for social ensembles, we could examine the encoding properties of an ensemble by examining the weights associated with that set of neurons in different classifiers. Specifically, in the EPM, it is unclear whether the open and closed arms are encoded by reciprocal changes in activity within the same neurons vs. by two distinct neuronal populations. E.g., it is possible that the neurons which increase their activity in the open arms decrease their activity in the closed arms and vice-versa so that open and closed arm representations are *anticorrelated*. Alternatively the set of neurons which increase or decrease their activity in the open arms may be distinct from those which do so in the closed arms so that open and closed arm representations are *orthogonal*. To distinguish between these possibilities, we use classifiers which compare activity in each arm (open or closed) to the baseline condition (rather than the other arm). As shown in Fig. 5B, in WT mice, weights for the closed arm ensemble in the open arm classifier cluster near zero. The same is true for weights for the open arm ensemble in the closed arm classifier. This suggests that open and closed arm ensembles are largely non-overlapping, such that representations of the open and closed arms are orthogonal, rather than anticorrelated.

**Figure 5.**
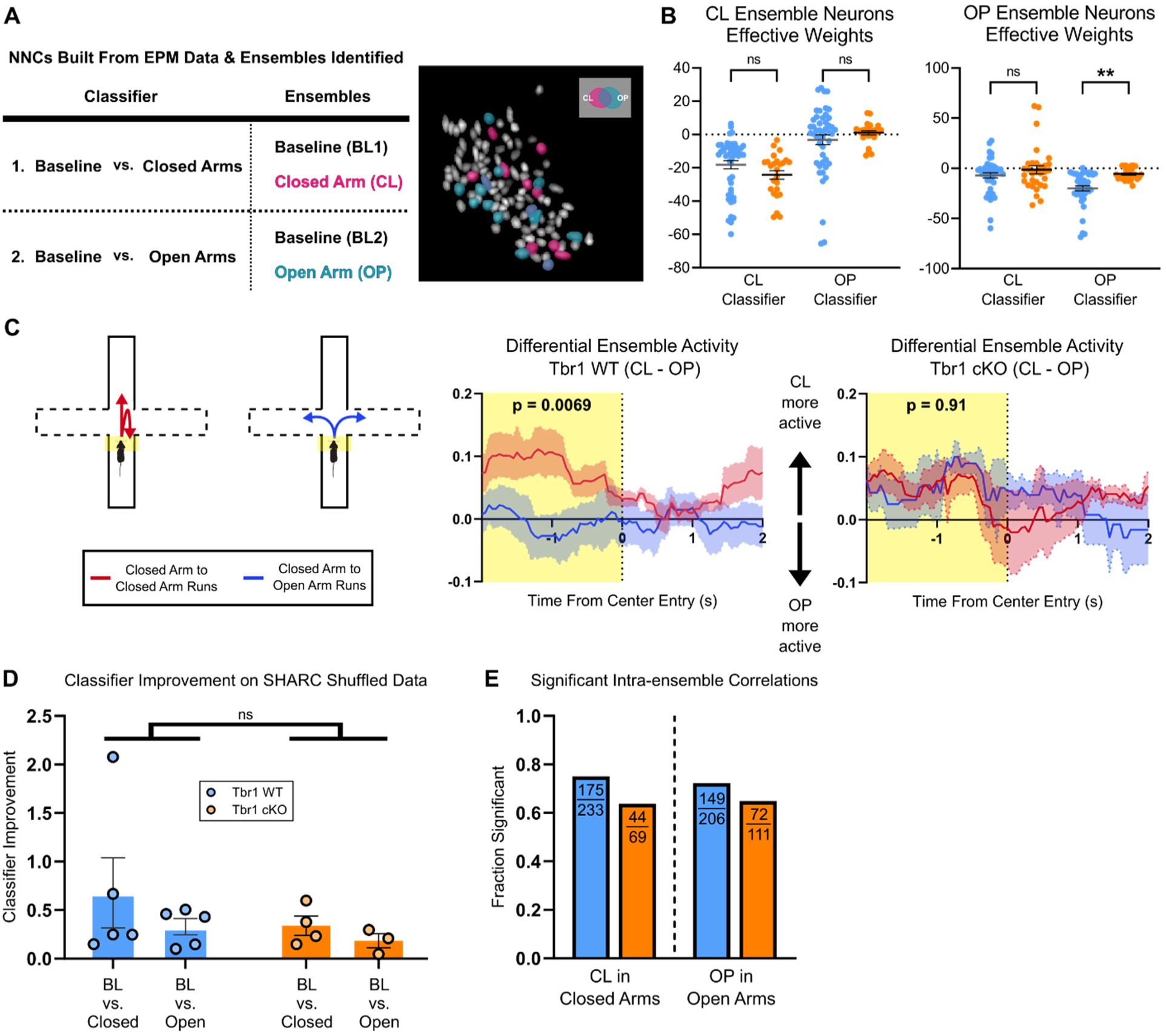
Encoding of the open arms of the EPM is disrupted in Tbr1 cKO mice. (A) *Left,* summary of NNCs trained on EPM data for each mouse and resulting ensembles, corresponding to neurons which encode the open (OP) or closed (CL) arms. *Right,* example FOV from one mouse with the two EPM ensembles highlighted. (B) *Left,* effective weights for neurons in the CL ensemble (n = 48 for WT, n = 25 for cKO) in NNCs for identifying periods of closed arm (left) or open arm (right) exploration. *Right,* effective weights of OP ensemble neurons (n = 43 for WT, n = 31 for cKO) in both open and closed arm NNCs. The magnitude of OP neuron weights in the OP classifier are significantly larger in WT than cKO mice, reflecting stronger encoding of the open arms. P-values shown are from Kruskal-Wallis tests with Dunn’s multiple comparison correction between genotypes. (C) Relative activity of CL and OP ensembles is predictive of arm transitions in WT, but not cKO mice. For each ensemble in each mouse, a normalized activity trace was calculated and aligned to Closed-Closed or Closed-Open arm transitions, and mean ensemble activity traces were computed. The Differential Ensemble Activity traces were calculated by subtracting the mean OP activity from the mean CL activity for each transition type, and then these Differential Activity traces were averaged across mice of the same genotype (*bottom left* for WT; *bottom right,* for cKO). A multi-factor ANOVA was then completed for each genotype to determine if the Differential Ensemble activity in the 2s before entry to the EPM center (regions highlighted in yellow) was significantly associated with the transition type. Transition type was revealed to be a significant factor in WT (p = 0.0069) but not cKO mice (p = 0.91). (D) Performance of NNCs trained on real data and tested on shuffled data. Performance of each trained classifier was assessed on Swap Shuffled and SHARC datasets, and a Performance Improvement score was calculated to assess the contribution of correlated activity in encoding EPM arms. A 2-way ANOVA with Sidaks correction of Performance Improvement scores showed no difference between genotypes (p = 0.322). (E) The fraction of significant pairwise correlations within each EPM ensemble during exploration of its corresponding maze arm is plotted. Pairwise correlations and their p-values were calculated using the Matlab *corr* function. Correlations with a p-value < 0.05 were taken as significant. P-values for differences in the fraction of of significant pairwise correlations between genotypes were computed for each ensemble using Fisher’s Exact tests.

Notably, effective weights for OP ensemble neurons in the OP Classifier were significantly weaker in cKO mice compared to WT (-19.9 ± 2.6 for WT vs. -5.7 ± 1.1 for cKO, Dunn’s post-hoc test, p = 0.0003), suggesting overall weaker encoding of the open arms in cKO mice (**Fig. 5B**). What might the consequences of this weaker encoding be? Many previous studies have used multineuron recordings to show that different sets of neurons, most notably in the mPFC and ventral hippocampus, increase their activity in the open or closed arms (Adhikari et al., 2011; A. T. Lee et al., 2019). However, it has been much more difficult to identify signals which predict future behavior, e.g., decisions to explore vs. avoid the open arms. Our previous study on prefrontal vasoactive intestinal polypeptide (VIP)-expressing interneurons was able to do this, but by using bulk activity from a specific interneuron subtype rather than multineuron patterns of activity (A. T. Lee et al., 2019). One reason it has been challenging to identify predictive signals from multineuron recordings is that the number of open arm entries in an individual mouse is relatively low, limiting the available data and making it impractical to, for example, train a classifier on approach vs. avoidance trajectories. We hypothesized that open and closed arm ensembles, derived from NNCs, might represent a useful tool for overcoming this challenge. Specifically, instead of training a classifier to identify ensembles specifically associated with approach vs. avoidance trajectories, we hypothesized that open and closed arm ensembles might themselves anticipate upcoming decisions. To test this hypothesis, we examined the activity of CL and OP ensembles time-locked to the moment of EPM center entry. For each animal, we identified all periods of center exploration lasting one second or longer, then classified these based on the arm (closed or open) from which the animal came, and into which the animal subsequently transited. This yielded four run types: closed-closed, closed-open, open-open, and open-closed.

Normalized ensemble activity traces were calculated for each CL and OP ensemble as follows: first, the binary calcium event rasters were summed across an ensemble, resulting in a time series representing how many neurons within an ensemble were active in each frame.

Next, each time series was normalized to its maximum value over the entire EPM recording. Finally, we used 4-second windows around each center zone entry to average together all time-locked segments of these time series corresponding to each run type (closed-closed, closed-open, open-closed, open-open). We first averaged time-locked time series for each run type within each mouse, then across all mice of the same genotype. This yielded 16 averaged ensemble activity traces that were all aligned to the time of center entry (2 ensemble types x 4 run types x 2 genotypes).

Inspecting these ensemble activity traces revealed a relationship between ensemble activity and approach-avoidance decisions in WT mice. During the 2s preceding center entry, CL ensemble activity was higher than OP ensemble activity, specifically for closed-closed runs but not for closed-open runs. In other words, stronger activity in the CL ensemble, relative to the OP ensemble, tended to predict open arm avoidance. However, this relationship was no longer evident in cKO mice. To more clearly illustrate this, we plotted the differential ensemble activity (CL-OP) for both run types (closed-closed vs. closed-open) (**Fig. 5C**). Specifically, the normalized OP activity vector was subtracted from the normalized CL activity vector for the entire EPM recording to yield the CL-OP differential activity vector for each animal. Then, the mean differential activity was calculated for each type of run for a 4-second window around EPM center entry. To assess the statistical validity of these observations, we performed a multi- factor ANOVA on mean CL-OP differential activity during the 2s preceding center entry for each genotype, using the run-type and animal ID as factors. Indeed, this revealed a significant effect of run-type for WT, but not *Tbr1* cKO mice (WT: Run Type main effect, p = 0.0069; Mouse ID main effect, p = 0.064; Run Type x Mouse ID interaction, p = 0.23; cKO: Run Type main effect, p = 0.91; Mouse ID main effect, p = 0.17; Run Type x Mouse ID interaction, p = 0.92).

### The Encoding of EPM behavior by correlated L5 activity is not altered in Tbr1 cKO mice

Given that correlated ensemble activity is disrupted during social behavior in cKO mice, we wondered whether a similar perturbation would occur during EPM exploration in cKO mice. As described previously for social behavior, NNCs trained on real data recorded during EPM exploration were tested using both SHARC and Swap shuffled datasets (which preserve or disrupt correlations, respectively). In this case, there was no significant difference between WT and cKO mice in the degree to which classifier performance was improved using SHARC instead of swap-shuffling (**Fig. 5D**). There was also no difference in the proportion of significant intra-ensemble PWCs across genotypes for either the CL or OP ensemble (**Fig. 5E**).

### Activity levels are no longer lower in Tbr1 cKO mPFC neurons after lithium treatment

A previous study of ours in *Tbr1* cKO mice found that a single dose of lithium chloride (LiCl) could reverse both synaptic and social behavior abnormalities in these animals (Fazel Darbandi et al., 2020, 2022). This raises the question of whether LiCl treatment would also normalize the encoding of socioemotional behaviors. To address this, we injected a subset of our mice with a single dose of LiCl (5 mg/kg) one day after initial EPM testing, then retested these animals 30 days later on both the serial interaction assay and the EPM (**Fig. 6A**). Of note, because the number of these animals was very limited, we were not able to include an additional cohort of mice which receive vehicle only. Thus, this experiment could only examine whether the abnormalities in neuronal activity patterns we initially observed are still present in cKO mice even after LiCl treatment, which is known to normalize abnormalities in social behavior and synapse number.

**Figure 6.**
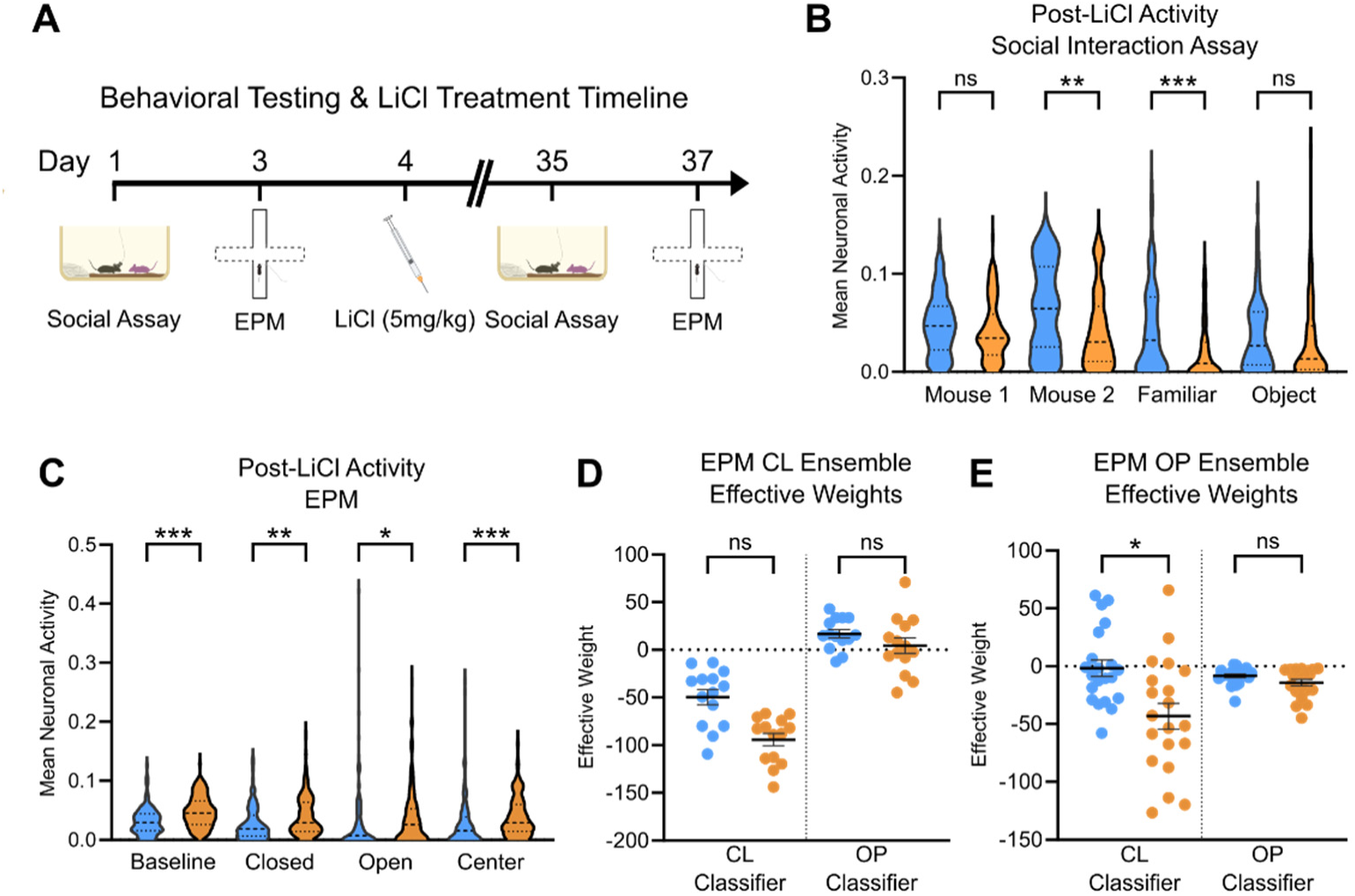
Lithium treatment rescues deficient activity in Tbr1 cKO neurons. (A) Experimental timeline for behavior testing and lithium treatment. All animals, WT and cKO, were given LiCl after initial social and EPM testing. Animals were then retested 30 days after lithium administration. (B) Violin plots of activity of all recorded neurons during the serial social interaction assay after lithium treatment (n=93 WT, n=98 cKO). Mean activity of each neuron was calculated as the fraction of frames corresponding to active social exploration that had a detected calcium event. P-values shown are from a Kruskal-Wallis test between genotypes. (C) Violin plots of activity of all recorded neurons during the EPM assay after lithium treatment (n=113 WT, n=118 cKO). Mean activity of each neuron was calculated as the fraction of frames corresponding to exploration of each EPM zone that had a detected calcium event. P-values shown are from a Kruskal-Wallis test between genotypes. (D) Effective weights of CL ensemble neurons (n = 14 for WT, n = 14 for cKO) from both trained EPM NNCs. P-values shown are from Kruskal-Wallis tests with Dunn’s multiple comparison correction between genotypes. (E) Effective weights of OP ensemble neurons (n = 21 for WT, n = 20 for cKO from both trained EPM NNCs. After LiCl treatment, the magnitude of OP neuron weights in the OP classifier are no longer significantly different between genotypes, and are in fact non-significantly larger in magnitude in cKO than WT (contrast with the smaller OP neuron weights in the OP classifier for cKO pre-treatment: Fig. 5B). P-values shown are from Kruskal-Wallis tests with Dunn’s multiple comparison correction between genotypes.

Whereas before LiCl treatment, the activity of *Tbr1* cKO neurons was deficient during all epochs of the serial interaction assay and in all zones of the EPM (**Fig. 1D-E**), after treatment, deficits were only seen during the Mouse2 and Familiar interactions (Mouse1: WT = 0.0481 ± 0.0033, cKO = 0.0422 ± 0.0031; Mouse2: WT = 0.0650 ± 0.0047, cKO = 0.0428 ± 0.0040; Familiar: WT = 0.0444 ± 0.0048, cKO = 0.0215 ± 0.0029; Object: WT = 0.0381 ± 0.0038, cKO = 0.0309 ± 0.0039; p = 0.999, 0.0031, 0.0003, 0.327, respectively) (**Fig. 6B**). In fact, after LiCl, *Tbr1* cKO neurons were actually more active than their WT counterparts in all zones of the EPM, as well as during the Baseline period immediately prior to the EPM (Baseline: WT = 0.0321 ± 0.0022, cKO = 0.0481 ± 0.0024; Closed Arms: WT = 0.0282 ± 0.0027, cKO = 0.0401 ± 0.0031; Open Arms: WT = 0.0305 ± 0.0056, cKO = 0.0375 ± 0.0044; Center: WT = 0.0287 ± 0.0041, cKO = 0.0393 ± 0.0031; p = 0.0007, 0.0051, 0.0201, 0.0002, respectively) (**Fig. 6C**).

Furthermore, after LiCl, we no longer observed a significant reduction in the magnitude of open arm ensemble (OP) classifier weights in the open arm (OP) classifier in cKO. In fact, after treatment these weights were non-significantly larger in magnitude for cKO neurons compared to WT (WT = -8.41 ± 1.56, cKO = -14.30 ± 2.89; p > 0.9999) (**Fig. 6E**), indicating that the pre-treatment deficit in open arm encoding was no longer present.

## DISCUSSION

There has been a persistent gap in understanding how disruptions in ASD-linked genes alter patterns of neuronal activity, leading to the behavioral abnormalities observed in mouse models. Identifying these changes in neural activity is critical for two reasons. First, this elucidates the pathophysiological chain of causation, linking the abnormalities in cells and synapses caused by genetic disruptions to changes in the patterns of activity which ultimately shape behavior. Second, behavioral measurements in mouse models are inherently superficial and lack access to the internal state of the mouse. By contrast, measurements of neural activity can confirm that abnormal behavior reflects changes in the processing of relevant behavioral information, distinguish between many possible information processing deficits which might plausibly lead to the same behavioral phenotype, and have greater sensitivity to confirm the presence of abnormalities when behavioral readouts are subtle or ambiguous.

In this context, our results reveal specific abnormalities in how the prefrontal cortex encodes information related to social interaction and approach-avoidance decisions in *Tbr1* cKO mice. In the process, we also identify several novel aspects of how the prefrontal cortex encodes socioemotional information. Finally, we additionally showed that treatment with lithium may reverse neural activity deficits, not just behavioral and synaptic abnormalities, in *Tbr1* cKO mice.

### Disrupted social encoding by correlations is a convergent mechanism in ASD models

*Tbr1*-cKO mice display reduced social interaction with a novel conspecific, as well as (under some conditions) decreased anxiety-related avoidance in the EPM. During these behaviors, mPFC layer 5 *Tbr1*-cKO neurons display a marked reduction in activity. The encoding of social interaction by neuronal coactivity is diminished, and within neuronal ensembles encoding social behavior, inter-neuronal correlations are disrupted. These results confirm a previous result from our lab that correlations between mPFC neurons play an important role in encoding social information, and parallel our specific finding that the ability of correlations to enhance social encoding is diminished in a different ASD mouse model: *Shank3* KO mice (Frost et al., 2021). Interestingly, mPFC neurons in the *Shank3* mice were hyperactive (relative to WT counterparts) during social interaction, whereas we observe decreased activity in *Tbr1*-cKO mice. Despite these divergent changes in activity levels, we find a similar circuit-level deficit in correlations, suggested that disruptions in correlated activity within the prefrontal cortex may represent a convergent mechanism contributing to social deficits across distinct ASD models and etiologies.

An interesting aspect of this finding is that in cKO mice, it is specifically the weak negative correlations between neurons in the same social ensemble which are lost. The presence of significant negative correlations within WT but not cKO social ensembles suggests that neurons within a social ensemble normally tile a social interaction and/or exhibit specificity for particular sub-behaviors, and that this type of tiling / specificity is lost in cKO mice.

One unifying framework for understanding how diverse genetic disruptions lead to ASD is the excitation-inhibition (or E/I) balance within mPFC circuits (Sohal & Rubenstein, 2019). E/I balance is hypothesized to play a crucial role in processing and encoding social information, as inhibition can reduce noise, potentially by coordinating the activity of local excitatory neurons. Previous work has demonstrated that mPFC E/I balance is crucial for normal social behavior (Yizhar et al., 2011), and that restoring normal E/I balance also restores normal social behavior in the *Cntnap2* mouse model of ASD (Selimbeyoglu et al., 2017). Further, another study established that this *Cntnap2* mouse model had reduced pairwise correlations between mPFC neurons, although this finding was not specifically in the context of social interaction (Levy et al., 2019). Together, these findings suggest that by perturbing E/I balance, disruptions in ASD risk-genes may compromise correlated patterns of activity that normally support social behavior.

### Prefrontal ensembles encode social interactions

We also identified distinct, but overlapping, ensembles of neurons in the mPFC that encode different types of social interactions. Specifically, the social interaction paradigm we used involved interactions with two different novel conspecifics, along with a repeat presentation of one of these conspecifics as a “familiar” interaction. This allowed us to evaluate the consistency of mPFC representations across different social interactions. We found that ensembles encoding the two different novel interactions were significantly non-overlapping in cKO animals, but not in WT, implying that representations of different conspecifics become abnormally distinct in the cKO. Furthermore, we find that neurons encoding the familiar mouse (which was a second exposure to Mouse 1), are similarly activated during Mouse 1 and Familiar interactions, but significantly less active during Mouse 2 interactions for both genotypes. These results show that mPFC neurons are tuned to conspecific identity, and that this encoding is not necessarily disrupted by the loss of *Tbr1*.

### Reactivation and preactivation of social ensembles

We also investigated the activity of neuronal ensembles encoding social interaction during the rest periods between successive interactions. Surprisingly, social ensembles in *Tbr1* WT mice displayed elevated activity during the rest period immediately following the first interaction with Mouse 1 (Figure 3). While the M1 ensemble neurons had the most striking rest period activity during HC2, elevated activity also occurred within the Familiar and M2 ensembles. For all social ensembles, this rest period activity was absent *Tbr1* cKO animals. To our knowledge, this is the first description of reactivation for neurons encoding social experiences in the prefrontal cortex, and the first time that the preactivation of social ensembles has been described in any structure.

One possible explanation for this reactivation (of the M1 ensemble) and “preactivation” (of the M2 and Familiar ensembles) is that the mPFC contains an intrinsic network of neurons that are primed for recruitment during social behavior, and that this entire network (not just the neurons encoding a specific interaction) is recruited following an initial social interaction.

Previous studies have demonstrated the general importance of mPFC for the formation of long term memories (DeNardo et al., 2019; Preston & Eichenbaum, 2013), including social memories. A recent study using fos-TRAP identified a population of L5 mPFC neurons that underly social memory (Xing et al., 2021). It is tempting to speculate that the absence of reactivation and preactivation in *Tbr1* cKO mice is related to the abnormally distinct M1 and Familiar representations observed in this model. Specifically, reactivation and preactivation may serve to link together representations for social experiences that are close together in time, related to the more general concept of ‘memory linking’ (Zaki et al., 2025).

One question is whether the absence of reactivation and preactivation we observe in *Tbr1* cKO mice represents a deficit that is specific for social circuitry vs. a disruption of processes related to replay and reactivation more generally. Related to this, the specific circuits which normally mediate social ensemble reactivation and preactivation, and whose malfunction gives rise to the deficits observed in *Tbr1* cKO mice, are unclear. On the one hand, since these mutant mice specifically lack *Tbr1* in L5 neocortical neurons it is tempting to hypothesize that these neurons play a key role in driving reactivation and preactivation. On the other hand, previous work has implicated hippocampal inputs to mPFC in long term social memory formation, and the acute reactivation of social memory neurons has been observed during sharp-wave ripples in the ventral CA1, suggesting a possible role for ventral CA1 inputs to mPFC in social reactivation (and possibly preactivation as well) (Tao et al., 2022). Notably, this previous study found that reactivation of ventral CA1 social neurons was disrupted in *Shank3* KO mice, suggesting that impaired social reactivation may be another conserved mechanism across etiologically distinct ASD models.

### Ensemble activity predicts approach-avoidance decisions

Interestingly, the contribution of coactivity to encoding of the open and closed arms of the EPM is unaffected in cKO animals, suggesting that different circuit mechanisms regulate prefrontal coactivity during social vs. approach-avoidance behaviors. However, we do find that overall encoding of the anxiogenic, open arms of the EPM is weakened, which may explain why these mice tend to explore the open arms more than WT mice. Furthermore, the activity of open and closed arm ensembles was predictive of approach-avoidance decisions in the center of the EPM in WT animals, but not in cKO mice.

A large body of work has shown that mPFC neurons differentially encode the open vs. closed arms of the EPM (Adhikari et al., 2011; Padilla-Coreano et al., 2016), that theta-frequency synchronization between the ventral hippocampus (vHPC) and mPFC changes as mice approach the center of the EPM (Adhikari et al., 2010; Cunniff et al., 2020; Jacinto et al., 2016; Padilla-Coreano et al., 2016), and that acute manipulations of the mPFC affect open arm exploration (Kjaerby et al., 2016; A. T. Lee et al., 2019; Padilla-Coreano et al., 2016). All of this suggests an important role for the prefrontal cortex in neural computations related to approach-avoidance decisions, but it has been difficult to identify populations of prefrontal neurons actively engaged in these decisions. In this context, our finding that prefrontal ensembles which encode the open vs. closed arms also predict decisions to approach vs. avoid the open arms ∼1-2 seconds before mice even enter the center zone, reveals specific neuronal ensembles involved in this process, and suggests that the differential activity of closed vs. open arm ensembles underlies this computation.

Interestingly, a recent study identified two populations of ventral hippocampus (vHPC) to mPFC projection neurons that influence approach-avoidance decisions, and which may thus provide input which drives the predictive prefrontal ensemble activity we found (Sánchez-Bellot et al., 2022). Given that previous studies of these *Tbr1*-cKO animals demonstrated a loss of synaptic input to L5 cKO neurons (Fazel Darbandi et al., 2020), the loss of predictive encoding in the cKO animals may reflect diminished input to mPFC from one or both of these vHPC populations. Interestingly, previous work from our laboratory found disruptions in vHPC-mPFC communication associated with decreases in avoidance of the EPM open arms in another mouse model of ASD (*Pogz*^+/-^ mice) (Cunniff et al., 2020). This suggests a possible mechanism whereby diminished vHPC-mPFC input causes the loss of predictive ensemble activity within mPFC, leading to reduced anxiety-related avoidance. This could represent yet another pathophysiological mechanism conserved across etiological distinct models of ASD. Future work could test this potential convergent mechanism, e.g., by examining whether vHPC-mPFC synaptic input is specifically affected by the loss of *Tbr1* and/or whether the loss of *Pogz* disrupts predictive signals within mPFC ensembles.

### Therapeutic implications

Finally, we examined the effects of lithium treatment on prefrontal activity in *Tbr1* cKO mice. Previously, a single dose of lithium was shown to reverse synaptic and behavioral abnormalities in these *Tbr1* cKO animals (Fazel Darbandi et al., 2020). While our lithium treatment cohort was underpowered to probe many changes in ensembles and correlations, this treatment does appear to reverse deficits in the activity levels of mPFC cKO neurons during social and approach-avoidance behaviors. It also restores normal open arm encoding in the EPM, as measured by ensemble weights in a neural network classifier. While a larger cohort of mice including saline-treated controls would be needed to confirm these findings and understand how lithium affects patterned mPFC activity, this provides promising preliminary evidence that lithium treatment may elicit a normalization of neural activity (and the processing of socioemotional information), not just a superficial improvement in gross behavioral measurements.

## ACKNOWLEDGEMENTS

This work was supported by the UCSF Dolby Family Center for Mood Disorders and the Simons Foundation Autism Research Initiative (SFARI grant 630332 to J.L.R.R. and SSDC grant 736613 to V.S.S.).

## AUTHOR CONTRIBUTIONS

M.L.T and V.S.S designed the experiments and analysis. S.R.S. performed the behavioral experiments for the large behavior-only cohort of mice. M.L.T. performed all other rodent experiments and data analysis. M.L.T. and V.S.S. wrote the manuscript. J.L.R.R. and S.F.D. provided important input on experimental design and the manuscript.

---

**Figure S1.**
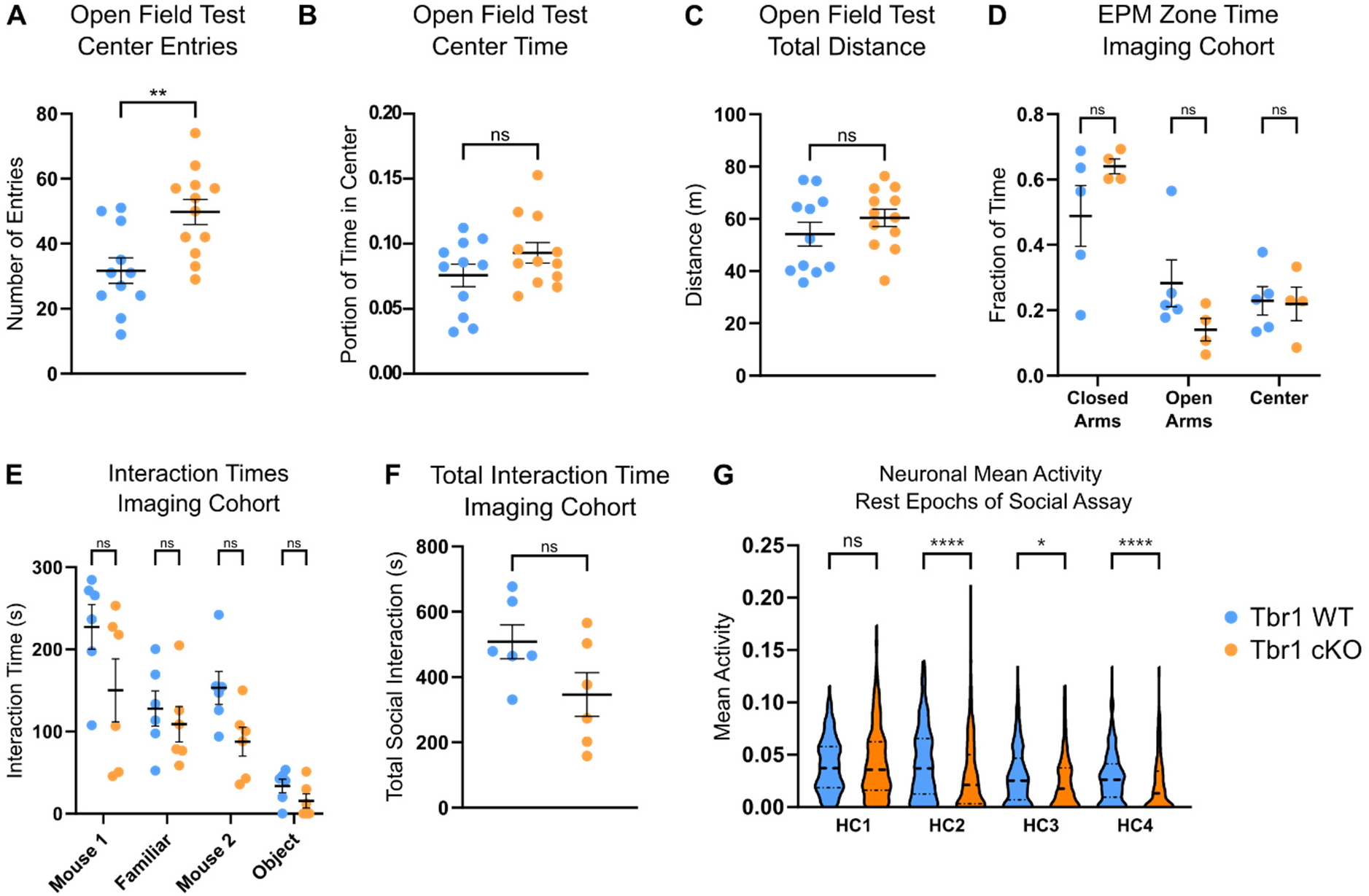
Related to Figure 1, additional quantifications of behavior and neural activity. (A) Center zone entries during a 10-minute Open Field Test (n= 23 mice: 11 WT, 12 Tbr1 cKO). Data is from the same behavioral animal cohort shown in Fig. 1A-B. P-value from a Welch’s Unpaired t-test. (B) Portion of time spent in center zone during OFT. P-value from a Welch’s Unpaired t-test. (C) Total distance traveled during the OFT. P-value from a Welch’s Unpaired t-test. (D) EPM zone time for the imaging cohort (n = 9 mice: 5 WT, 4 Tbr1 cKO). P-values are from a 2-way ANOVA with Sidak’s multiple comparison test. (E) Interaction times during the serial interaction assay for the imaging cohort of mice (n = 6 WT and 6 Tbr1 cKO mice). P-values are from a 2-way ANOVA with Sidak’s multiple comparison test. (F) Total social interaction time (Mouse, Familiar, and Mouse 2 combined) for the imaging cohort. P-value from a Welch’s Unpaired t-test. (G) Violin plots showing the distribution of mean activity for mPFC L5 neurons during rest epochs of the serial interaction assay (n=306 WT, n=278 cKO). Mean activity of each neuron was calculated as the fraction of frames during each rest epoch that had a detected calcium event. P-values shown are from a Kruskal-Wallis test between genotypes.

**Figure S2.**
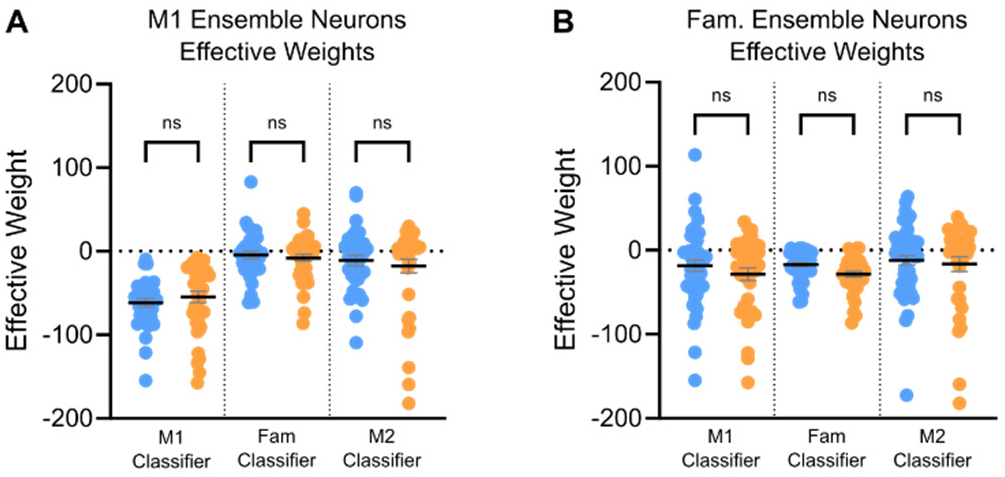
Related to Figure 2, classifier weights for the M1 and Familiar ensembles. (A) Effective weights of neurons in the M1 ensemble (n = 35 and 40 for WT and cKO, resp.) in each of the three trained NNCs from (Fig. 2B). P-values from Kruskal-Wallis tests with Dunn’s multiple comparison correction between genotypes. (B) Effective weights of neurons in the Familiar (Fam) ensemble (n = 48 and 36 for WT and cKO, resp.) in each of the three trained NNCs from (Fig. 2B). P-values from Kruskal-Wallis tests with Dunn’s multiple comparison correction between genotypes.

**Figure S3.**
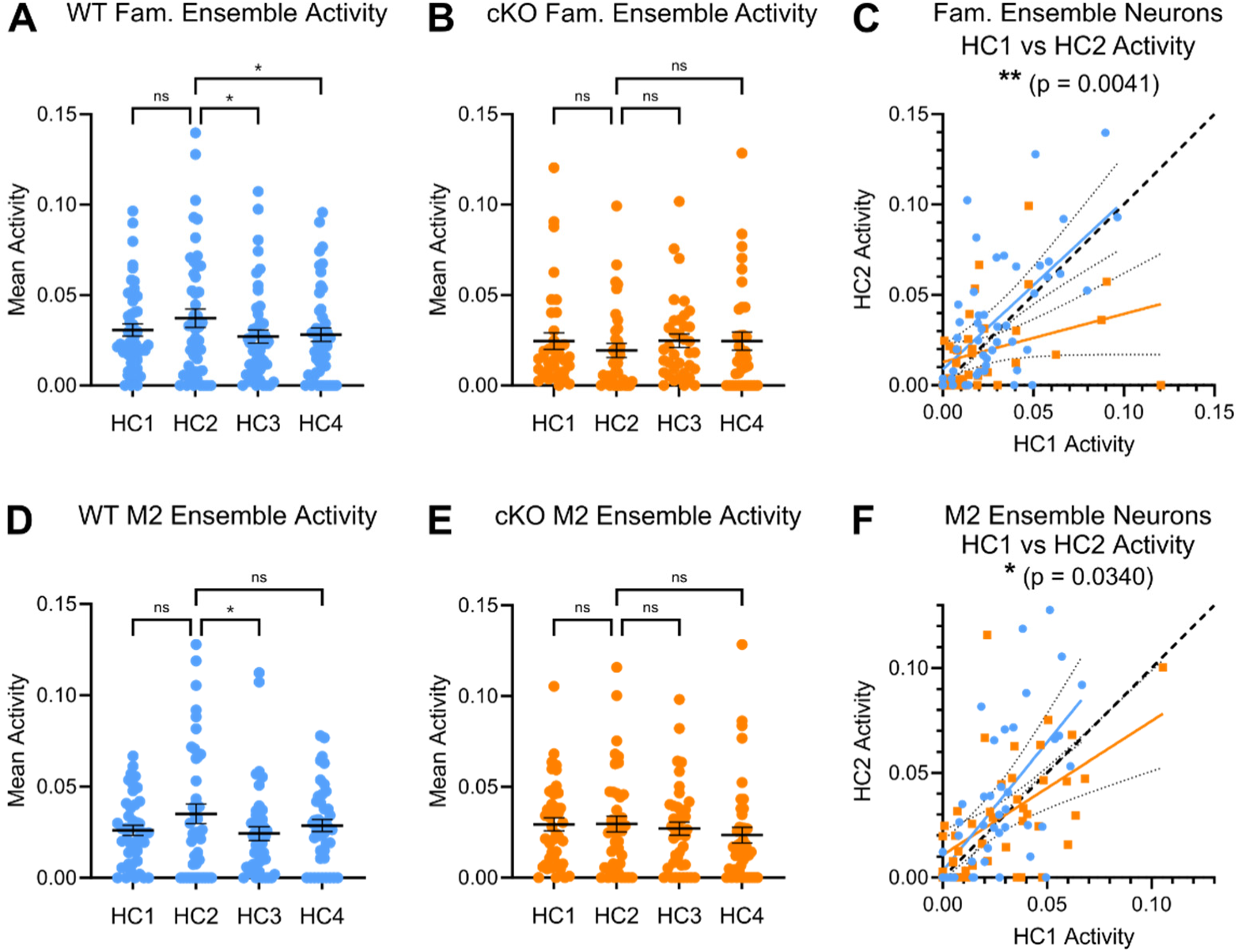
Related to Figure 3, activity of neurons in the Familiar and M2 ensembles across various home cage epochs. (A) Activity of WT neurons in the Familiar (Fam) ensemble (n = 48) during rest epochs of the serial interaction assay. P-values are from a RM one-way ANOVA with Dunnett’s multiple comparison test, comparing activity during each rest epoch vs. during the HC2 epoch. (B) Activity of cKO neurons in the Fam ensemble (n = 36) during rest epochs of the serial interaction assay. P-values are from a RM one-way ANOVA with Dunnett’s multiple comparison test, comparing activity during each rest epoch vs. during the HC2 epoch. (C) Scatter plot displaying the relationship between HC1 and HC2 activity of WT (blue) and cKO (orange) Fam neurons. The best fit lines from simple linear regression analyses have significantly different slopes, indicating that WT Fam neurons display stronger activation during this epoch. (D) Activity of WT neurons in the M2 ensemble (n = 43) during rest epochs of the serial interaction assay. P-values are from a RM one-way ANOVA with Dunnett’s multiple comparison test, comparing activity during each rest epoch vs. during the HC2 epoch. (E) Activity of cKO neurons in the M2 ensemble (n = 42) during rest epochs of the serial interaction assay. P-values are from a RM one-way ANOVA with Dunnett’s multiple comparison test, comparing activity during each rest epoch vs. during the HC2 epoch. (F) Scatter plot displaying the relationship between HC1 and HC2 activity of WT (blue) and cKO (orange) M2 neurons. The best fit lines from simple linear regression analyses have significantly different slopes, indicating that WT M2 neurons display stronger activation during this epoch.

**Figure S4.**
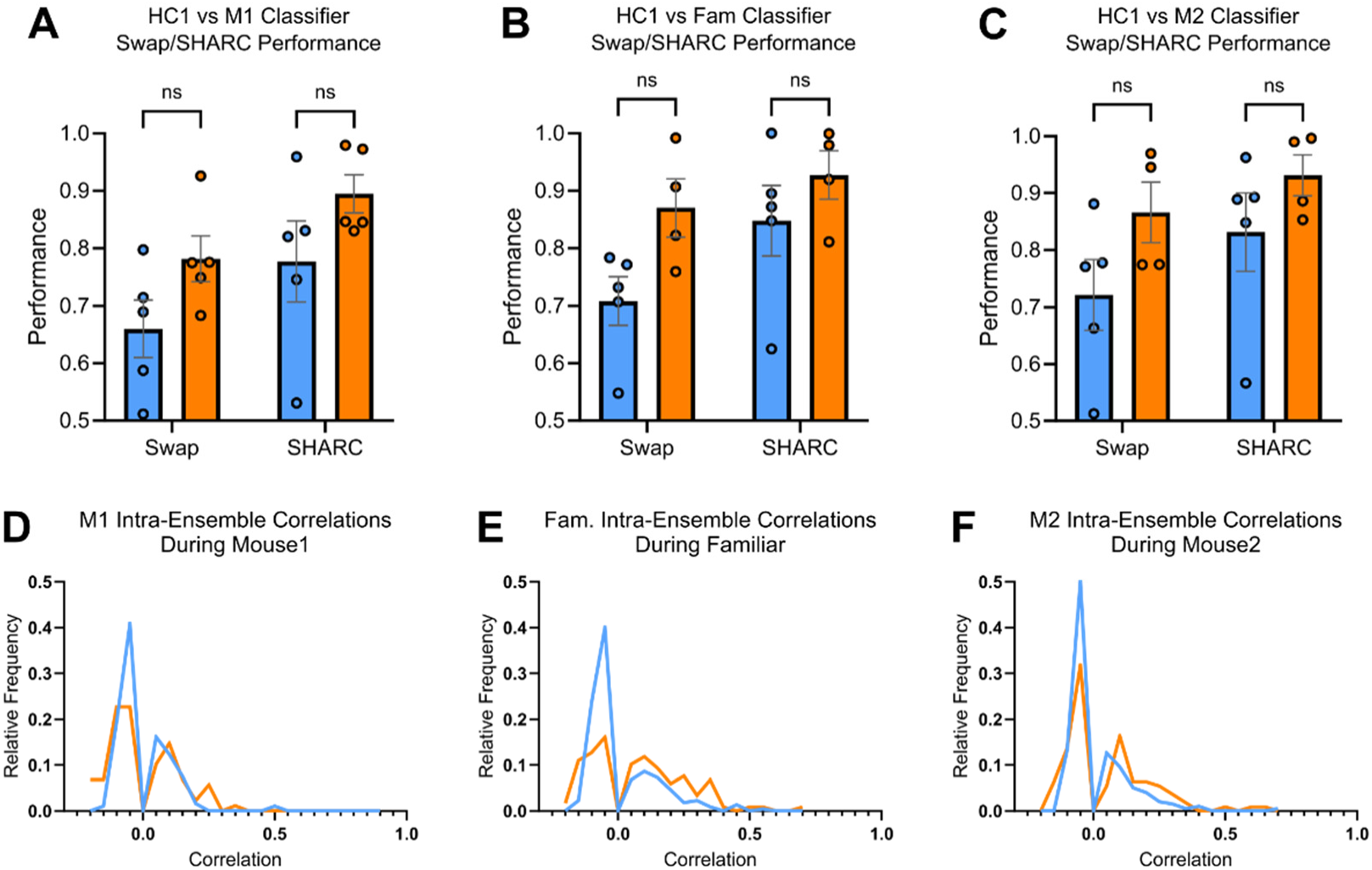
Related to Figure 4, classifier performance and distributions of significant intra-ensemble correlations. (A) Performance of M1 classifiers on Swap and SHARC shuffled data used to calculate the “Performance Improvement” plotted in Fig. 4B (n = 10; 5 WT and 5 cKO mice). P-values shown are from a 2-way ANOVA with Sidak’s multiple comparisons test. (B) Performance of Familiar (Fam) classifiers on Swap and SHARC shuffled data used to calculate the “Performance Improvement” plotted in Fig. 4B (n = 5 WT and 4 cKO mice; 1 cKO animal was excluded due to insufficient interaction time with the Familiar conspecific). P-values are from 2-way ANOVA with Sidak’s multiple comparisons test. (C) Performance of M2 classifiers on Swap and SHARC shuffled data used to calculate the “Performance Improvement” plotted in Fig. 4B (n = 5 WT and 4 cKO mice; 1 cKO animal was excluded due to insufficient interaction time with the Mouse 2 conspecific). P-values are from a 2-way ANOVA with Sidak’s multiple comparisons test. (D) Distributions of significant intra-ensemble pairwise correlations of neurons in the M1 ensemble during Mouse 1 interaction, corresponding to Fig. 4C. (E) Distributions of significant intra-ensemble pairwise correlations of neurons in the Fam ensemble during Familiar interaction, corresponding to Fig. 4C. (F) Distributions of significant intra-ensemble pairwise correlations of neurons in the M2 ensemble during Mouse 2 interaction, corresponding to Fig. 4C.

**Figure S5.**
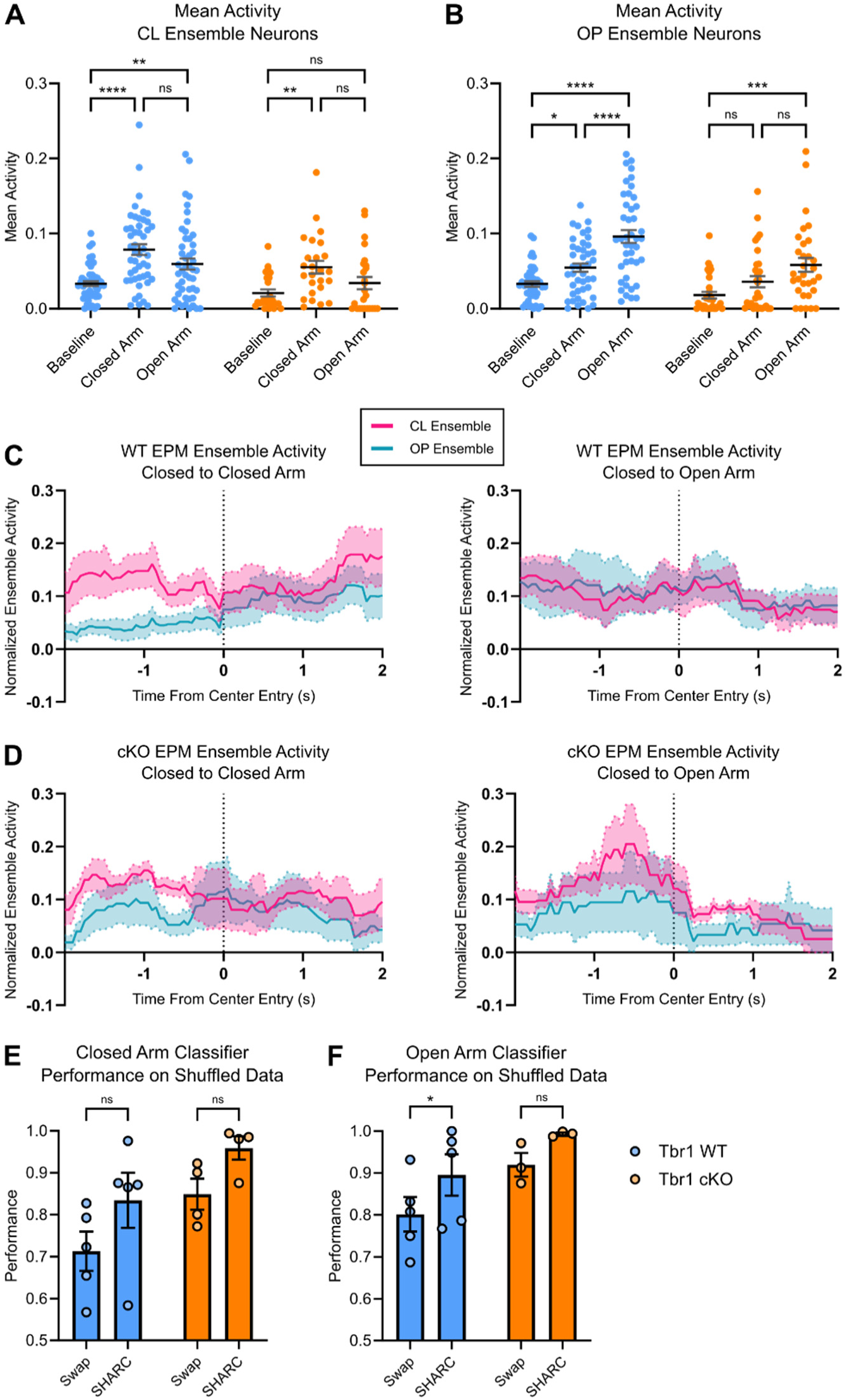
Related to Figure 5, EPM ensemble activity and classifier performance. (A) Mean activity of CL ensemble neurons during the Baseline period and in different zones of the EPM (n = 48 WT neurons, 25 cKO neurons). P-values are from 2-way ANOVA with Tukey’s multiple comparison test. (B) Mean activity of OP ensemble neurons during the Baseline period and in different zones of the EPM (n = 43 WT neurons, 31 cKO neurons). P-values are from 2-way ANOVA with Tukey’s multiple comparison test. (C) Normalized activity traces of WT CL and OP ensembles aligned to EPM center entry for Closed-to-Closed (left) and Closed-to-Open (right) transitions. These traces were used to calculate the Differential Ensemble Activity traces shown in Fig. 5C. (D) Same as (C), but for cKO CL and OP ensembles. (E) Performance of Closed Arm classifiers on Swap and SHARC shuffled data used to calculate the “Performance Improvement” plotted in Fig. 5D (n = 5 WT and 4 cKO mice). P-values are from a 2-way ANOVA with Sidak’s multiple comparisons test. (F) Performance of Open Arm classifiers on Swap and SHARC shuffled data used to calculate the “Performance Improvement” plotted in Fig. 5D (n = 5 WT and 3 cKO mice; 1 cKO animal was excluded due to insufficient time spent in the open arms). P-values are from a 2-way ANOVA with Sidak’s multiple comparisons test.

**Figure S6.**
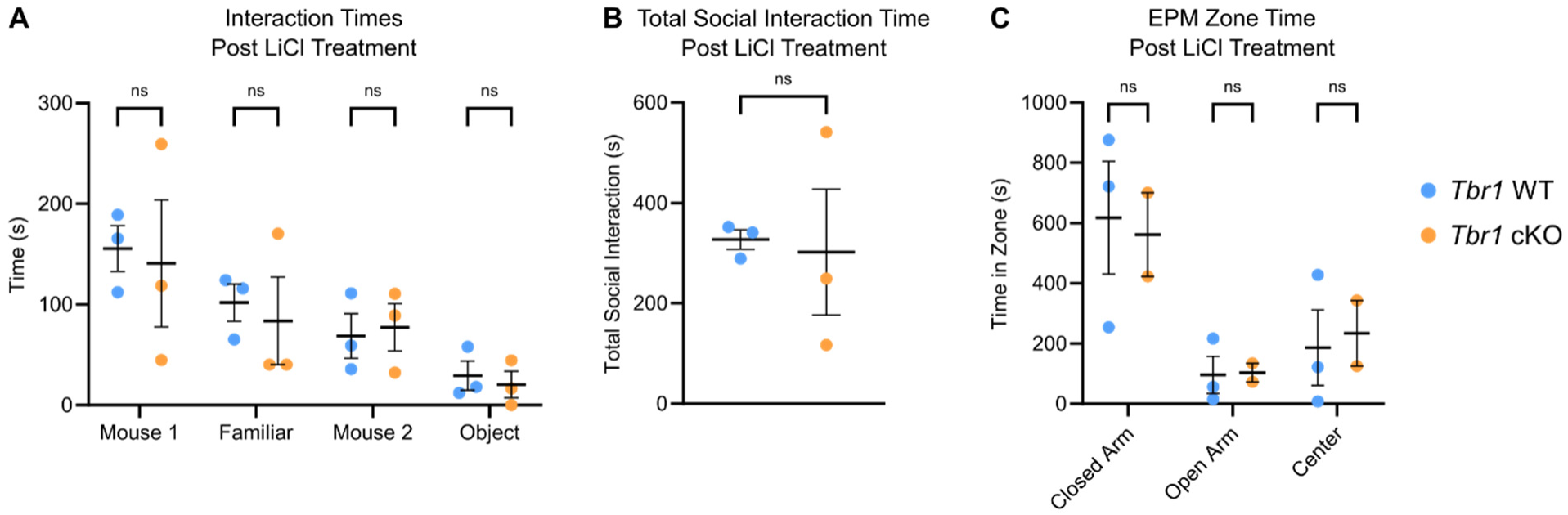
Related to Figure 6, post-LiCl treatment animal behavior. (A) Interaction times during the serial interaction assay following LiCl treatment (n = 6 mice: 3 WT, 3 Tbr1 cKO). P-values from 2-way ANOVA with Sidak’s multiple comparison test. (B) Total social interaction time (Mouse, Familiar, and Mouse 2 combined) following LiCl treatment. P-value from a Welch’s Unpaired t-test. (C) EPM zone time from a 15-minute EPM test following LiCl treatment (n = 5 mice: 3 WT, 2 Tbr1 cKO). P-values from 2-way ANOVA with Sidak’s multiple comparison test.

